# Dynamic competition between SARS-CoV-2 NSP1 and mRNA on the human ribosome inhibits translation initiation

**DOI:** 10.1101/2020.08.20.259770

**Authors:** Christopher P. Lapointe, Rosslyn Grosely, Alex G. Johnson, Jinfan Wang, Israel S. Fernández, Joseph D. Puglisi

## Abstract

SARS-CoV-2 recently emerged as a human pathogen and is the causative agent of the COVID-19 pandemic. A molecular framework of how the virus manipulates host cellular machinery to facilitate infection remains unclear. Here, we focus on SARS-CoV-2 NSP1, which is proposed to be a virulence factor that inhibits protein synthesis by directly binding the human ribosome. Using extract-based and reconstitution experiments, we demonstrate that NSP1 inhibits translation initiation on model human and SARS-CoV-2 mRNAs. NSP1 also specifically binds to the small (40S) ribosomal subunit, which is required for translation inhibition. Using single-molecule fluorescence assays to monitor NSP1–40S subunit binding in real time, we demonstrate that eukaryotic translation initiation factors (eIFs) modulate the interaction: NSP1 rapidly and stably associates with most ribosomal pre-initiation complexes in the absence of mRNA, with particular enhancement and inhibition by eIF1 and eIF3j, respectively. Using model mRNAs and an inter-ribosomal-subunit FRET signal, we elucidate that NSP1 competes with RNA segments downstream of the start codon to bind the 40S subunit and that the protein is unable to associate rapidly with 80S ribosomes assembled on an mRNA. Collectively, our findings support a model where NSP1 associates with the open head conformation of the 40S subunit to inhibit an early step of translation, by preventing accommodation of mRNA within the entry channel.

**SIGNIFICANCE STATEMENT:** SARS-CoV-2 is the causative agent of the COVID-19 pandemic. A molecular framework for how SARS-CoV-2 manipulates host cellular machinery to facilitate infection is needed. Here, we integrate biochemical and single-molecule strategies to reveal molecular insight into how NSP1 from SARS-CoV-2 inhibits translation initiation. NSP1 directly binds to the small (40S) subunit of the human ribosome, which is modulated by human initiation factors. Further, NSP1 and mRNA compete with each other to bind the ribosome. Our findings suggest that the presence of NSP1 on the small ribosomal subunit prevents proper accommodation of the mRNA. How this competition disrupts the many steps of translation initiation is an important target for future studies.

## INTRODUCTION

Beta-coronaviruses (CoVs) are a family of RNA viruses that include human pathogens^1^. In the last two decades, two CoVs have emerged from animal hosts to cause epidemic diseases of the human respiratory tract: Severe Acute Respiratory Syndrome (SARS-CoV, in 2002)^2,3^ and Middle East Respiratory Syndrome (MERS-CoV, in 2012)^4^. A third CoV emerged in late 2019 — SARS-CoV-2 — that is responsible for the ongoing COVID-19 pandemic^5^. Given the lack of effective therapies against SARS-CoV-2, there is an urgent need for a molecular understanding of how the virus manipulates the machineries present in human cells.

SARS-CoV-2 and the closely-related SARS-CoV have single-stranded, positive-sense RNA genomes nearly 30 kb in length^6,7^. Upon entry of a virion into human cells, the genomic RNA is released into the cytoplasm where it must hijack human translation machinery to synthesize viral proteins^8^. As the genomic RNA has a 7-methylguanosine (m^7^G) cap on the 5’-terminus, viral protein synthesis likely proceeds via a process reminiscent of that which occurs on typical human messenger RNAs (mRNAs)^9^. However, as viral proteins accumulate, human translation is inhibited and host mRNAs are destabilized, which facilitates suppression of the host immune response^10–13^.

Studies on SARS-CoV have implicated non-structural protein 1 (NSP1), the first encoded viral protein, as a virulence factor with a key role in the shutdown of host translation^10,11,14^. In infected cells or upon its ectopic expression, NSP1 inhibits human translation, which is dependent on its association with the small (40S) subunit of the human ribosome^12–17^. In a connected but separable activity, NSP1 destabilizes at least a subset of human mRNAs, likely via recruitment of an unidentified human endonuclease^12,13,15,16,18^. NSP1 from SARS-CoV-2 is expected to employ similar mechanisms, given its approximately 85% sequence identity with the SARS-CoV protein. Thus, NSP1 has a near singular ability to dramatically disrupt host gene expression; yet, the mechanism by which this inhibition occurs is not clear.

The 40S subunit is the nexus for translation initiation, recruiting an m^7^G-capped mRNA through a multistep, eukaryotic initiation factor(eIF)-mediated process. Prior to recruitment of an mRNA, the 40S subunit is bound by numerous eIFs, including eIF1, eIF1A, eIF3, eIF5, and the eIF2–Met-tRNA^Met^_i_–GTP ternary complex (TC)^19^. The eIFs make extensive contacts with the 40S subunit, including the ribosomal A and P sites^20,21^. They also manipulate the dynamics of the 40S head region to facilitate mRNA recruitment, which has structural consequences at both the mRNA entry (3’ side of mRNA) and exit (5’ end of mRNA) channels. Following mRNA recruitment and scanning of the 5’ untranslated region (UTR), a series of compositional and conformational changes occur^22,23^. This ultimately repositions the 40S subunit head into the closed conformation and accommodates the anticodon-stem-loop of the initiator tRNA at the start codon^20–23^. Taken together, the intrinsic dynamics of translation initiation present many opportunities and obstacles for NSP1 association with the ribosome, and its subsequent inhibition of translation.

Here, we merge biochemical and single-molecule approaches to probe the molecular function of SARS-CoV-2 NSP1 and its interaction with the human ribosome. We illustrated that NSP1 inhibited translation initiation on human and SARS-CoV-2 model mRNAs, and then determined how the NSP1– 40S subunit interaction was modulated by eIFs and mRNA. Our results reveal allosteric control of NSP1 association by eIF1 and likely eIF3j, and competition between NSP1 and mRNA to bind the ribosome. Combined with recent structural studies, our study suggests a mechanism for how NSP1 inhibits translation initiation.

## RESULTS

### NSP1 inhibited translation initiation of host and SARS-CoV-2 model mRNAs

The sequence conservation (≈85% identity) of SARS-CoV-2 NSP1 with the homologous protein from SARS-CoV suggested that it would inhibit human translation (**Sup. Fig. 1A**). To test this hypothesis, we employed a cell-free *in vitro* translation (IVT) assay using HeLa cellular extract. As a model of a host mRNA, the 5’ and 3’ UTRs from human GAPDH mRNA were fused to a nanoLuciferase (nLuc) open-reading frame (ORF) (**Fig. 1A**). This GAPDH reporter mRNA was added to IVT reactions that contained increasing concentrations of purified SARS-CoV-2 NSP1 (**Sup. Fig. 1B**). Following incubation for 45 minutes, we observed a concentration-dependent reduction in nLuc activity by wild-type NSP1 (**Fig. 1B**), with an *IC*_*50*_ of 510 ± 20 nM (95% C.I.; *R*^*2*^ =0. 83). Given its multi-functional potential, we also tested two mutant NSP1 proteins with alanine substitutions that replaced both the conserved RK124-125 residues (RK/AA) implicated in mRNA destabilization or the conserved KH164-165 residues (KH/AA) implicated in ribosome binding (**Sup. Fig. 1**). As predicted based on the SARS-CoV protein^16^, the NSP1(RK/AA) mutant inhibited translation similar to the wild-type protein (*IC*_*50*_ ≈ 420 ± 11 nM, 95% C.I., *R*^*2*^ = 0.89), while the NSP1(KH/AA) mutant failed to inhibit translation (**Fig. 1B**). The inhibitory effect of NSP1 on protein synthesis therefore may be meditated by an NSP1-ribosome interaction, independent of mRNA degradation.

**Figure 1.**
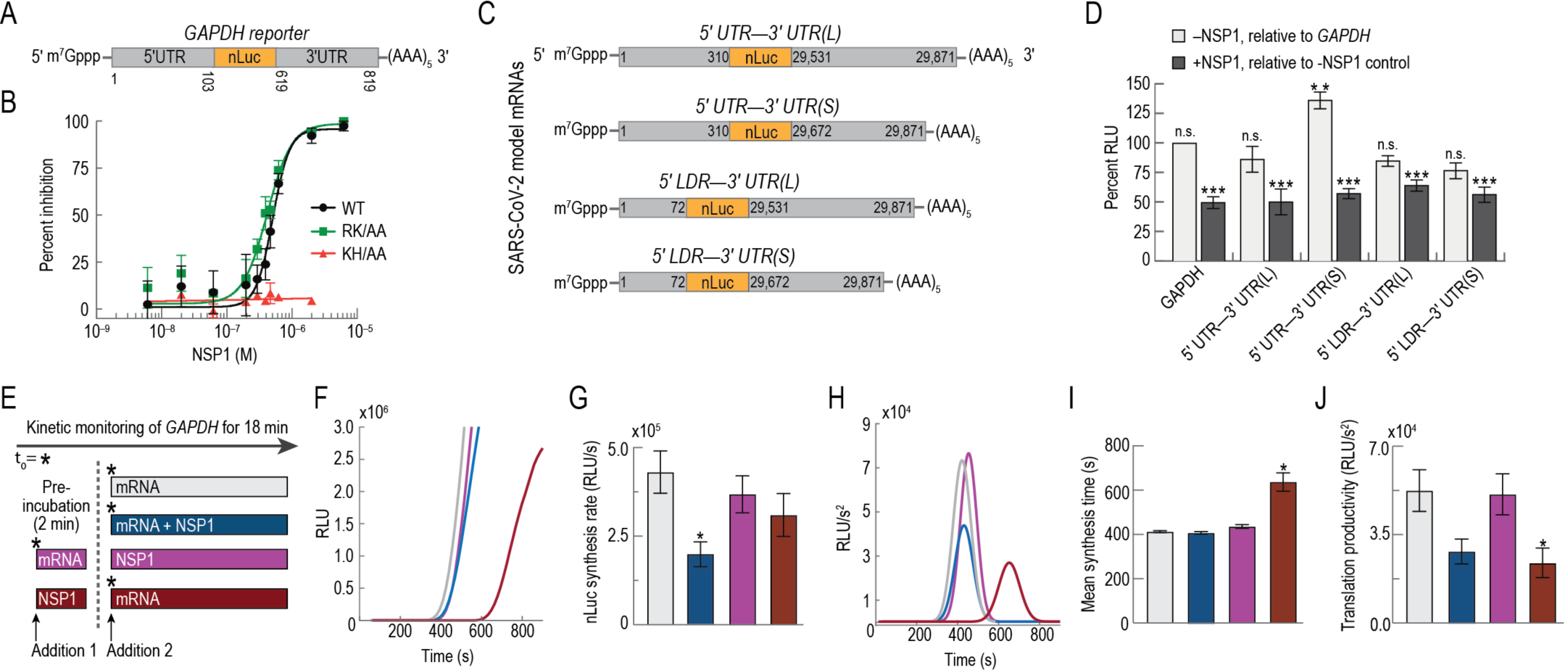
NSP1 from SARS-CoV-2 inhibited translation initiation. **A)**. Schematic of host model mRNA used in HeLa cell-free *in vitro* translation (IVT) assays. The nano-Luciferase (nLuc) coding sequence of the mRNA was flanked by the 5’ and 3’ UTRs of human GAPDH. Numbers refer to the nucleotide position in human GAPDH mRNA (NCBI GenBank accession: AF261085). **B)**. NSP1 dose-response analysis of GAPDH reporter mRNA IVT in HeLa extract treated with either wild-type (WT; *n*=3), a predicted ribosome-binding deficient mutant (KH/AA; n=2), or an RNA cleavage deficient mutant (RK/AA; n=2) NSP1. The mean response ± SEM (symbols, error bars) and curve fits (lines) from non-linear regression analysis of the data are plotted. WT *IC*_*50*_ = 510 ± 20 nM (95% C.I.; *R*^*2*^ =0. 83) and RK/AA *IC*_*50*_ = 420 ± 11 nM (95% C.I.; *R*^*2*^ = 0.89). **C)**. Schematic of SARS-CoV-2 nLuc reporter mRNAs used in HeLa cell-free *in vitro* translation (IVT) assays. The nLuc coding sequence was flanked by either the full-length viral 5’UTR or the subgenomic 5’ leader sequence (LDR) and either the 3’UTR(L), which contains part of N ORF or 3’UTR(S), which begins after ORF10. Numbers refer to nucleotide position in the viral genome (NCBI GenBank accession: MN997409.1). **D)**. Plot of the mean nLuc signal from cell-free translation of host and viral reporter mRNAs in the absence and presence (400 nM) of wild-type NSP1. Without NSP1 (light gray), mean translational activities (percent RLU) were compared to GAPDH reporter mRNA in the absence of NSP1 (** = *p* ≤ 0.0006; and n.s. = *p* ≥ 0.2, one-way ANOVA). GAPDH (*n*=6), 5’UTR-3’UTR(S) mRNA (*n*=3), 5’UTR-3’UTR(L) (*n*=3), and 5’LDR reporter mRNAs (*n*=5). In the presence of NSP1 (dark gray), samples were compared a control reaction that lacked NSP1 (*** = *p* ≤ 0.0008, t-test). GAPDH (*n*=6), viral 5’UTR-mRNAs (*n*=3), and viral 5’LDR-mRNAs (*n*=2). Error bars represent SEM. **E)**. Time-of-addition cell-free IVT experimental design. GAPDH reporter mRNA and WT NSP1 (400 nM) were added to HeLa IVT reactions in the order depicted above. The nLuc signal was continuously monitored *in situ*. Using the same color scheme as here, panels F-J depict results from six independent replicates for each experimental condition, except for the ‘preincubation with NSP1 reaction’ (red bar in E), which has *n*=3. **F)**. Time course of nLuc synthesis from a representative time-of-addition experiment. **G)**. Plot of mean synthesis rates determined from the plateau value of the first derivative of nLuc synthesis time course data. Mean values are plotted. Error bars represent SEM (\**p* = 0.02, one-way ANOVA). **H)**. Representative Gaussian fits of the second derivative of nLuc synthesis time course data shown in panel F. **I)**. Plot of mean synthesis time. Error bars represent SEM, * *= p* < 0.0001). **J)**. Plot of translational productivity. Error bars represent SEM, * *p* = 0.045, one-way ANOVA).

In the context of infection, full-length SARS-CoV-2 genomic RNA and its sub-genomic mRNAs must be translated in the presence of NSP1 protein. To examine whether RNA elements within the viral UTRs facilitate evasion of NSP1-mediated inhibition, we constructed model SARS-CoV-2 mRNAs in which the nLuc ORF was flanked on the 5’ end by either the full-length viral 5’UTR or the sub-genomic 5’ leader sequence (LDR) (**Fig. 1C**). At the 3’ end, we fused two different versions of the 3’UTR, beginning after the stop codon for N protein (L) or ORF10 (S), to account for ambiguity in ORF10 coding potential^24^. Translation of all four model viral mRNAs was reduced significantly by approximately 50% upon addition of 400 nM NSP1 relative to reactions that lacked NSP1 (*p* ≤ 0.0008, unpaired t-test) (**Fig. 1D**). The degree to which NSP1 inhibited translation of the viral reporters was similar to the inhibition observed for GAPDH reporter mRNA (*p* ≥ 0.2, one-way ANOVA). However, translation of the 5’UTR-3’UTR(S) model viral mRNA was 36% higher than that of the host and other viral reporters (*p* ≤ 0.0006, one-way ANOVA) in our IVT system (**Fig. 1D**). This may suggest that enhanced translational activity of viral RNAs relative to host mRNAs may play a role in SARS-CoV-2 infection.

To determine the phase of protein synthesis inhibited by NSP1, we employed a real-time translation assay in which nLuc activity was continuously monitored *in situ*. From the resulting time course data, we extracted the protein synthesis rate, mean synthesis time, and translational productivity^25^ from samples where NSP1 (400 nM) was either omitted from the IVT reaction, added simultaneously with mRNA (GAPDH), added after the reaction was pre-incubated with mRNA, or pre-incubated in the extract prior to mRNA addition (**Fig. 1E**). Pre-incubation of the extract with mRNA allows translation to initiate in the absence of NSP1, while pre-incubation with NSP1 primes the extract for inhibition. If NSP1 disrupts translation initiation, the inhibition would be dependent on the temporal availability of NSP1; otherwise, if NSP1 inhibits translation elongation or termination, the inhibition would be insensitive to the timing of NSP1 addition.

The impact of NSP1 on translation was dependent on its time of addition to the IVT reaction. Consistent with the end-point assays, we observed an approximate 2-fold reduction in the protein synthesis rate when NSP1 and GAPDH reporter mRNA were added simultaneously (*p* < 0.02, one-way ANOVA) (**Fig 1F,G**). While protein synthesis rates for the other samples were similar to the reaction that lacked NSP1, we observed a dramatic delay in the appearance of nLuc signal when cell extracts were pre-incubated with NSP1, but not vice versa (**Fig 1F**). This lag between the addition of mRNA (*t* = 0) and the initial appearance of nLuc signal is the time needed for a full round of translation (‘synthesis time’; sum of initiation, elongation, and termination). To compare synthesis times quantitatively, we fit the second derivative of the nLuc time course data to a Gaussian distribution (**Fig 1H**). In this analysis, the mean of the distribution represents the mean synthesis time and its amplitude is a gauge of translational productivity. As suggested by the raw data, the mean synthesis time when extracts were pre-incubated with NSP1 (637 ± 41 s) increased by 54% (p<0.0001, one-way ANOVA) compared to the reaction without NSP1 (413 ± 5 s), while mean times of the two other conditions (411 ± 6 s and 435 ± 8 s) were similar to the control (**Fig. 1I**). Pre-incubation with NSP1 also reduced translational productivity approximately 2-fold (*p* = 0.04, one-way ANOVA) (**Fig. 1J**). In contrast, pre-incubation with mRNA yielded mean synthesis times and translation productivity similar to reactions that lacked NSP1 (**Fig. 1I**,**J**). Thus, ribosomes pre-loaded with mRNA evaded NSP1-mediated inhibition, which strongly suggests that NSP1 is a potent inhibitor of translation initiation, perhaps linked to mRNA recruitment or accommodation.

### NSP1 stably associated with ribosomal pre-initiation complexes

To test whether NSP1 binds directly to the human 40S ribosomal subunit, we employed native gel shift assays with purified components. Using an 11-amino acid ybbR tag, single cyanine dye fluorophores were conjugated site-specifically onto purified NSP1 (**Supp.Fig. 2A,B**)^26,27^. When incubated with increasing concentrations of purified 40S subunits, the amount of fluorescently-labeled NSP1 that co-migrated with 40S subunits increased (**Supp. Fig. 2C**). In contrast, NSP1 did not co-migrate with human 60S or yeast 40S subunits (**Supp. Fig. 2D**). Whereas NSP1(KH/AA) was unable to block the NSP1–40S subunit interaction, inclusion of either wild-type NSP1 or NSP1(RK/AA) at 150-fold molar excess prevented co-migration of labeled NSP1 with human 40S subunits (**Supp. Fig. 2E**). Thus, NSP1 specifically binds to the human 40S ribosomal subunit, dependent on the presence of an intact KH motif in the C-terminus of the protein. Together with our extract-based assays, these data indicate that NSP1 binds the human 40S subunit to inhibit translation initiation.

**Figure 2.**
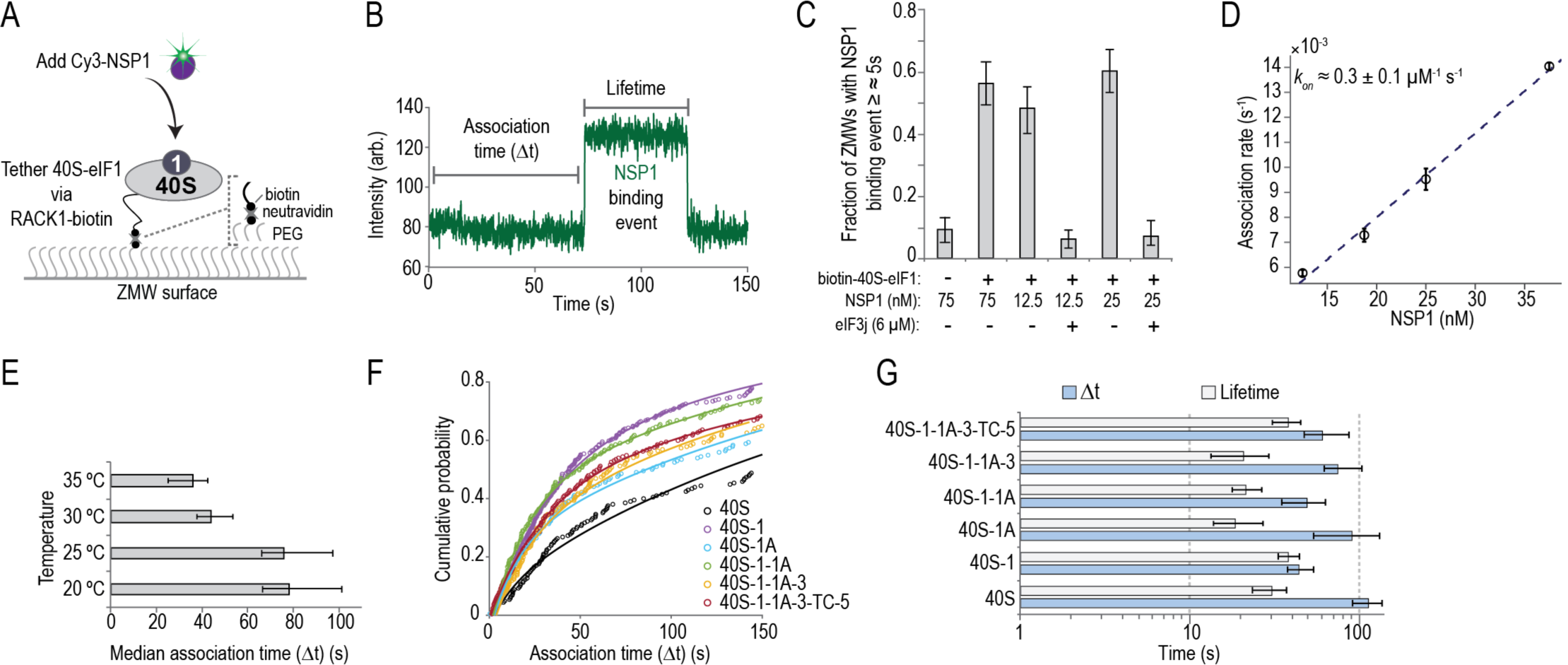
NSP1 associated with 40S subunits and most ribosomal pre-initiation complexes. **A)**. Experimental setup. Using a zero-mode-waveguide (ZMW) system, 40S ribosomal subunits biotinylated on RACK1 were tethered to a neutravidin-coated imaging surface within thousands of individual ZMWs in the presence of various eIFs (also see, **Supp. Figure 4B**). Upon start of data acquisition, Cy3-NSP1 (N-terminal ybbR tag) was added and fluorescence intensities were monitored. Association of Cy3-NSP1 with 40S subunits was detected via emission of Cy3 fluorescence. **B)**. Example single-molecule fluorescence trace that depicts association of Cy3-NSP1 with a tethered eIF1–40S subunit complex. Prior to tethering, 40S subunits were incubated with 30-fold molar excess eIF1. During imaging, eIF1 was present at 1 µM. The association time (Δt) was defined as the time elapsed from the addition of Cy3-NSP1 until the burst of Cy3 fluorescence (green), which signified NSP1 association. The lifetime was defined as the duration of the Cy3 fluorescence signal. **C)**. Plot of the fraction of ZMWs that contained at least one Cy3-NSP1 binding event ≥ ≈ 5s in duration in the indicated conditions at 20 °C. Error bars represent 99% confidence intervals (C.I.). **D)**. Plot of apparent association rates (open circles) of Cy3-NSP1 with tethered eIF1–40S subunit complexes at the indicated NSP1 concentrations at 20 °C. The dashed line represents a fit from linear regression analysis (adjusted *R*^2^ = 0.99), with a slope of 0.3 ± 0.1 and y-intercept of 0.0013 ± 0.002 (errors represent 95% C.I.). Error bars on the open circles represent 95% C.I. of the derived rates. **E)**. Plot of median association times of 25 nM Cy3-NSP1 (final concentration) with eIF1–40S subunit complexes at the indicated temperatures. Error bars represent 95% C.I. of the median values. **F)**. Plot of the cumulative probability of Cy3-NSP1 association times with the indicated ribosomal pre-initiation complexes. In all experiments, Cy3-NSP1 was present at 25 nM (final concentration) and the temperature was 30 °C. eIFs were pre-incubated with 40S subunits, and they were included at molar excess relative to 40S subunits during tethering and imaging to promote formation of the indicated complexes. eIF1, eIF1A, and eIF5 were at 1 µM; eIF2-GMPPNP-Met-tRNA^Met^_i_ (TC) at 100 nM; and eIF3 at 50 nM. Lines represent fits to double-exponential functions. See **Supp. Table 2** for samples sizes and parameters for fits. **G)**. Plot of median association times (light blue) and lifetimes (light gray) of Cy3-NSP1 with the indicated ribosomal pre-initiation complexes. Error bars represent 95% C.I. of the median.

Throughout translation initiation, there is a complex choreography between eIFs and the 40S ribosomal subunit. Yet, it was unknown how the NSP1–40S subunit interaction is affected by eIFs, either through direct interactions or induced changes in ribosome conformation. We therefore initially examined NSP1 binding to the 40S subunit in the presence of either 6 µM eIF1, eIF1A, or eIF3j. Each of these canonical eIFs bind with high affinity to the 40S subunit^28^ and may alter its conformation^20,21^. Inclusion of eIF1 increased the intensity of the NSP1–40S subunit band approximately 2-fold (mean ≈ 2 ± 0.4, 95% C.I.), while eIF3j eliminated the band (**Supp. Fig. 3A-C**). Unlike eIF1 and eIF3j, eIF1A had little impact on the NSP1–40S subunit complex (**Supp. Fig. 3A,D**). The NSP1–40S subunit interaction therefore was modulated inversely by two eIFs.

To define the kinetics of NSP1 binding to 40S subunits, we established a single-molecule assay to monitor NSP1 association with ribosomal pre-initiation complexes directly in real time. First, biotin was attached to purified 40S subunits that contained the ybbR tag on the ribosomal protein RACK1 (**Supp. Fig. 4A**), using the same strategy as with fluorescent dyes^29^. We then tethered preassembled eIF1–biotinylated-40S subunit complexes to thousands of zero-mode waveguide (ZMW) surfaces coated with neutravidin (**Supp. Fig. 4B**)^30^. Upon start of data acquisition, Cy3-NSP1 was added to the ribosomal complex (**Fig. 2A**), which had translation-inhibition and ribosome-binding activities similar to the wild-type protein (**Supp. Fig. 4C,D**). Association of NSP1 with the 40S subunit was manifested by a burst of Cy3 fluorescence (**Fig. 2B**). When NSP1 was added at 75 nM, the majority of ZMWs (56 ± 7 %) with tethered eIF1–40S subunit complexes contained at least one NSP1 binding event (≥ ≈ 5 s in length) (**Fig. 2C**). In contrast, the number of ZMWs with binding events was reduced dramatically in the absence of the tethered complex (9 ± 4 %). Similarly, at two different NSP1 concentrations, pre-incubation with 2.5 µM eIF3j reduced NSP1 binding to baseline levels (from 48 ± 7 % and 60 ± 7 %, to 6 ± 3 and 7 ± 3 %). Results consistent with specific binding also were obtained using total internal reflection fluorescence microscopy (TIRFM) at equilibrium (**Supp. Fig. 4E,F**). Thus, our assay directly monitored real-time association of NSP1 with tethered 40S ribosomal complexes and demonstrated competition by eIF3j for NSP1– 40S complex formation.

NSP1 bound the eIF1–40S subunit complex with high affinity. As predicted for a bimolecular interaction, NSP1 association times (the time elapsed from its addition until appearance of Cy3 signal) decreased with increasing concentration of NSP1 at 20 °C (**Fig. 2D** and **Supp. Fig. 4G**). Linear-regression analysis of the observed rates at various NSP1 concentrations yielded a bimolecular association rate of 0.3 ± 0.1 µM^-1^ s^-1^ (95% C.I.) (**Fig. 2D** and **Supp. Table 2**). The observed lifetime of the NSP1–40S subunit interaction (the duration of the Cy3 signal) was dependent on the power of the excitation laser (**Supp. Fig. 4H,I**), which indicated our measurements were limited by dye photostability. Nevertheless, with our longest measured lifetime as a lower bound (238 ± 6 s, 95% C.I.), we estimated that the equilibrium dissociation constant (*K*_*D*_) of the NSP1 interaction with eIF1–40S subunit complexes was ≤ 12.5 nM at 20 °C, similar to that of eIFs^28^.

NSP1 rapidly and stably associated with different ribosomal pre-initiation complexes. When added at 25 nM to eIF1–40S subunit complexes, median NSP1 association times (see, *Material and Methods*) were similar at 20 and 25 °C (67–101 s and 66–97 s, 95% C.I.), but they decreased nearly two-fold at 30 and 35 °C (38–54 s and 25–42 s, 95% C.I.) (**Fig. 2E** and **Supp. Fig. 4J**). We therefore measured NSP1 association times and lifetimes with 40S subunits in complex with various canonical eIFs at 30 °C. Consistent with our gel-based assays, the median NSP1 association time (at 25 nM) was decreased about 2-fold in the presence of eIF1 relative to 40S subunits alone (38–54 s versus 91–137 s, 95% C.I.) (**Fig. 2F**,**G** and **Supp. Fig. 4K**). Further inclusion of eIF1A, eIF3 that lacked the 3j subunit (eIF3), eIF5, and/or an eIF2–Met-tRNA^Met^_i_ –GMPPNP ternary complex (TC-GMPPNP) also yielded modest reductions in NSP1 association times. NSP1 lifetimes on the various eIF–40S subunit complexes were similar (**Fig. 2G** and **Supp. Fig. 4L**) and likely limited by dye photostability. Together with the temperature-dependence, the eIF-mediated modulation of NSP1 association with the 40S subunit, particularly by eIF1 and eIF3j, suggested that NSP1 may associate with a particular conformation of the 40S ribosomal subunit.

### mRNA within the entry channel of the 40S subunit inhibited NSP1 association

While this work was in progress, multiple groups reported structures of NSP1 bound to the human ribosome^31,32^. Despite its apparent flexibility, the N-terminal globular domain of NSP1 was localized to the solvent-exposed surface of the 40S subunit, near the entrance to the mRNA entry channel (**Supp.Fig. 5A**). This domain appears anchored by the two most C-terminal *α*-helices of NSP1, which were dynamic and unstructured in the free SARS-CoV NSP1 structure solved by NMR^33^; in the NSP1–40S subunit complex, these helices were well-resolved, docked within the mRNA entry channel where they contact ribosomal proteins uS3 and uS5, and helix 18 of the 18S rRNA. Further guided by our findings above, we therefore hypothesized that association of NSP1 with the 40S subunit would be sensitive to the conformation of the mRNA entry channel and to the presence of mRNA within it.

To test our hypothesis, we used the internal ribosome entry site (IRES) from hepatitis C virus (HCV), a structured RNA that directly binds to the human 40S subunit with high affinity (2–4 nM)^34^ (**Supp.Fig. 5B**). A flexible segment of the IRES (domain II) swivels the head of the 40S subunit to open the mRNA entry channel^35–37^ and allows accommodation of the mRNA coding region downstream from the start codon. We generated five HCV IRES model RNAs that were 5’-biotinylatyed and contained 0, 6, 12, 24, or 48 nucleotides downstream (3’) of the start codon (**Fig. 3A**). Based on the above structural models, mRNAs with more than 6 nucleotides after the start codon are predicted to at least partially occlude the NSP1 binding site within the mRNA entry channel. Following incubation of the biotinylated RNAs with fluorescently-labeled (Cy5 dye) ribosomal subunits, IRES–40S subunit complexes were tethered to a ZMW surface, and Cy3-NSP1 was added at 25 nM upon start of data acquisition (**Fig. 3B,C**).

**Figure 3.**
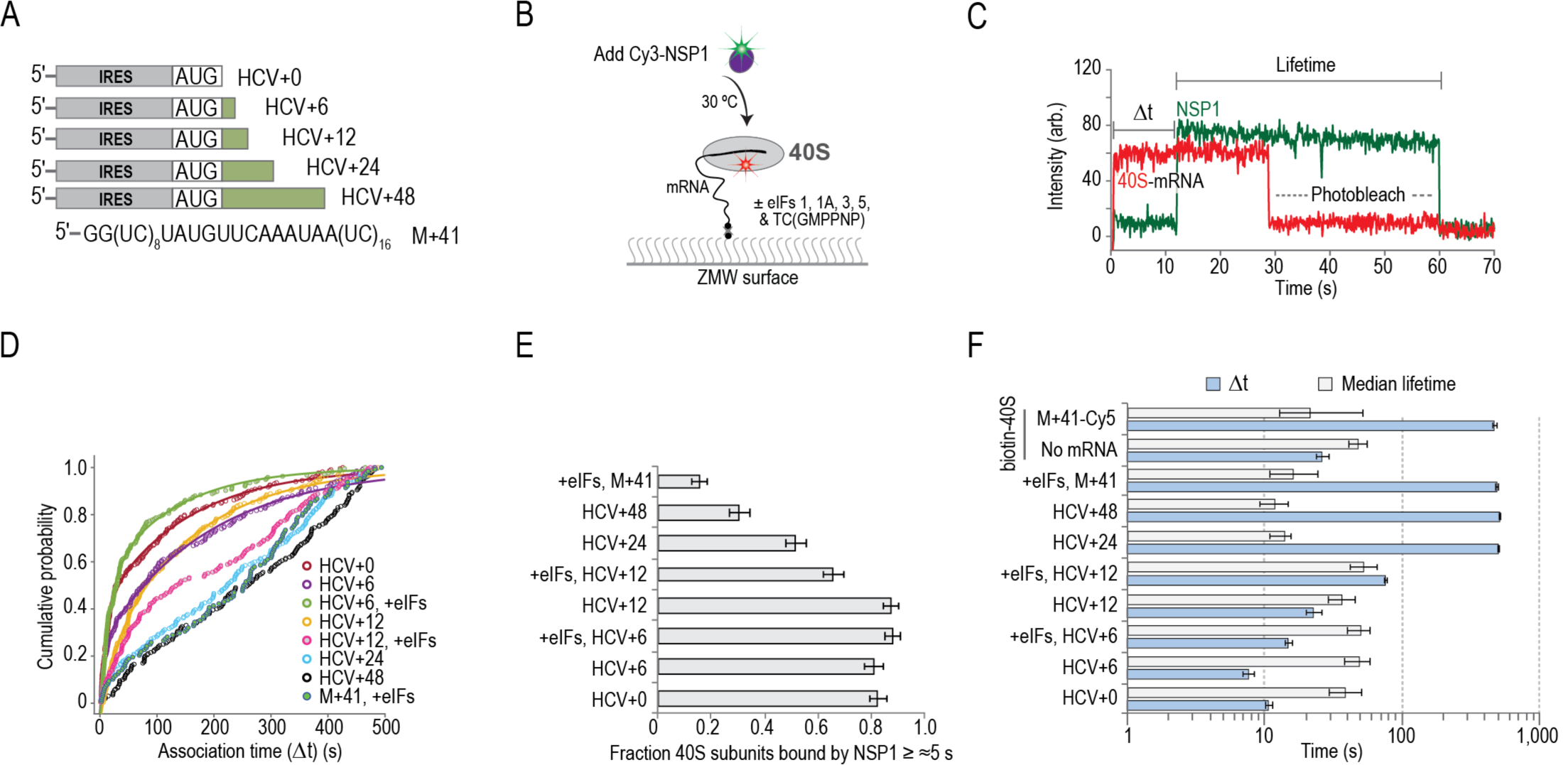
mRNA within the mRNA entry channel of the 40S subunit inhibited NSP1 association. **A)**. Schematic of model mRNAs. HCV IRES RNAs were biotinylated on the 5’-terminus and contained 0, 6, 12, 24, or 48 nucleotides downstream of the start codon (AUG). The model mRNA (M+41) contained 41 nucleotides downstream of the start codon, and it was biotinylated or labeled with Cy5 on the 3’-terminus. **B)**. Schematic of the single-molecule fluorescence experimental setup. 40S ribosomal subunits were labeled with Cy5 dye via RACK1-ybbR. Pre-formed mRNA–40S-Cy5 complexes (± indicated eIFs) were tethered to the ZMW imaging surface. At the start of data acquisition, Cy3-NSP1 (N-terminal ybbR tag) was added at 25 nM (final concentration) at 30 °C. **C)**. Example single-molecule fluorescence trace that depicts a tethered 40S-HCV+0 complex and subsequent association of NSP1. The 40S subunit and ybbR-NSP1 were labeled with Cy5 (red) and Cy3 (green) dyes, respectively. Loss of fluorescence signal due to dye photobleaching is indicated. Raw fluorescence intensities were corrected in this image to set baseline intensities to zero for presentation. The association time (Δt) was defined as time elapsed from the addition of Cy3-NSP1 until the burst of Cy3 fluorescence (green), which signified NSP1 association. The lifetime was defined as the duration of the Cy3 fluorescence signal. **D)**. Plot of the cumulative probability of observed Cy3-NSP1 association times with the indicated mRNA–40S subunit complexes at 30 °C. Cy3-NSP1 was added at 25 nM (final concentration) in all experiments. ‘+eIFs’ indicates eIFs 1, 1A, 3, 5, and TC(GMPPNP) were included at all stages of the experiment. Lines represent fits to single-or double-exponential functions. See **Supp. Table 3** for samples sizes and parameters for fits. **E)**. Plot of the fraction of the indicated mRNA–40S subunit complexes bound at least once by NSP1 for ≥ ≈ 5s. Error bars represent 99% C.I.. **F)**. Plot of the apparent association times (light blue) or median lifetimes (light gray) of NSP1 association with the indicated mRNA–40S subunit complexes. Apparent association times were defined as the reciprocal of the fast association rate derived from fits to double-exponential or linear functions, with error bars indicating the 95% C.I. of the rate. Error bars for median lifetimes represent 95% C.I.. See **Supp. Table 3** for samples sizes and parameters for fits.

Short and long segments of RNA downstream of the start codon had opposite effects on NSP1 association. With HCV+0 and HCV+6, NSP1 efficiently (80 ± 3% and 81 ± 3%, 95% C.I.) and rapidly associated (apparent *k*_*on*_^*-1*^ ≈ 11 ± 0.6 s and 8 ± 0.7 s, 95% C.I.) with tethered IRES-40S subunit complexes when added at 25 nM (**Fig. 3D-F** and **Supp. Table 3**). NSP1 association to this complex was approximately 2.5-fold faster than to the eIF1– 40S subunit complex (apparent *k*_*on*_^*-1*^ ≈ 29 ± 0.2 s, 95% C.I.) (**Fig. 3F** and **Supp. Fig. 5C**). In contrast, NSP1 associated less efficiently (52 ± 4% and 31 ± 4%) and much more slowly (apparent *k*_*on*_^*-1*^ ≈ ≥ 520 s) with HCV+24 and HCV+48 complexes relative to HCV+0 (**Fig. 3D-F**). We reasoned that the relative lack of inhibition we observed on HCV+12 could be due to inefficient accommodation of the mRNA into the entry channel. Indeed, inclusion of eIFs utilized for initiation by the HCV IRES (eIF1, eIF1A, eIF5, eIF3, and TC-GMPPNP) exacerbated the increase in NSP1 association times on HCV+12 to nearly 8-fold slower (apparent *k*_*on*_^*-1*^ ≈ 77 ± 2 s, 95% C.I.) relative to HCV+0 (**Fig. 3D-F**). In parallel to inhibited association, we also observed at least 3-fold decreased median NSP1 lifetimes on HCV+24 and HCV+48 complexes (11–16 s and 9–15 s, 95% C.I.) relative to the dye-limited measurements on HCV+0 and HCV+6 (29–50 s and 38–58 s) (**Fig. 3F** and **Supp. Fig. 5D**). We therefore used the same strategy as above to estimate that the lifetime of NSP1 on the 40S– HCV+0 complex was ≥ ≈230 s (**Supp. Fig. 5E**). Consequently, the *K*_*D*_ of the NSP1 interaction with the IRES–40S subunit complex was increased at least 500-fold (from ≤ ≈4 nM to ≥ ≈2–3 µM) by long segments of RNA downstream of the start codon.

To test whether our findings were generalizable to other RNAs, we preformed analogous experiments as above in two formats using an unstructured model mRNA (M+41) that contained 41 nucleotides downstream of the start codon (**Fig. 3A**). In the first, 3’-biotinylated M+41 RNA bound to 40S-Cy5 subunits were tethered to the imaging surface (**Supp.Fig. 5F,G**). In the second, 40S-biotin subunits bound to fluorescently-labeled M+41 were tethered (**Supp.Fig. 5H,I**). In both scenarios, we observed inefficient association of NSP1 with the mRNA–40S subunit complexes (16 ± 3% with 40S-biotin, 95% C.I.) and at least 48-fold increases in NSP1 association times (apparent *k*_*on*_^*-1*^ ≈ ≥ 500 s for both) relative to HCV+0 or tethered ribosomal complexes that lacked the model mRNA (apparent *k*_*on*_^*-1*^ ≈ 27 ± 3 s, 95% C.I.) (**Fig. 3D-F, Supp.Fig. 5J,K**, and **Supp. Table 3**). NSP1 association with mRNA–40S subunit complexes therefore was inhibited markedly by RNA segments downstream of the start codon that at least partially occlude its binding site in the mRNA entry channel.

Intriguingly, NSP1 also has been visualized bound to 80S ribosomal complexes^31^. To examine whether NSP1 could associate with 80S ribosomes, we used CRISPR-Cas9 and homology-directed repair to establish a FRET signal between the 40S and 60S subunits of the ribosome, analogous to our signal in yeast^38^. The ybbR tag was appended to all endogenous copies of ribosomal proteins uS19 (40S subunit) or uL18 (60S subunit) (**Supp.Fig. 6A-D**), which are within predicted FRET distance (≈50 Å) in structural models of 80S ribosomes (**Fig. 4A**). The tagged ribosomes were functional in cells (**Supp.Fig. 6E**), and purified 40S-ybbR and 60S-ybbR subunits were labeled efficiently (50– 80%) with Cy3 (FRET donor) and Cy5 (FRET acceptor) fluorescent dyes, respectively (**Supp.Fig. 6F**). After incubation with the IRES from the intergenic region of cricket paralysis virus (CrPV IRES), which assembles ribosomal subunits into 80S ribosomes independent of eIFs^39^, we observed a FRET efficiency distribution (mean ≈ 0.5 ± 0.01, 95% C.I.) between the labeled 40S and 60S subunits, consistent with structural predictions (**Fig. 4B,C**).

**Figure 4.**
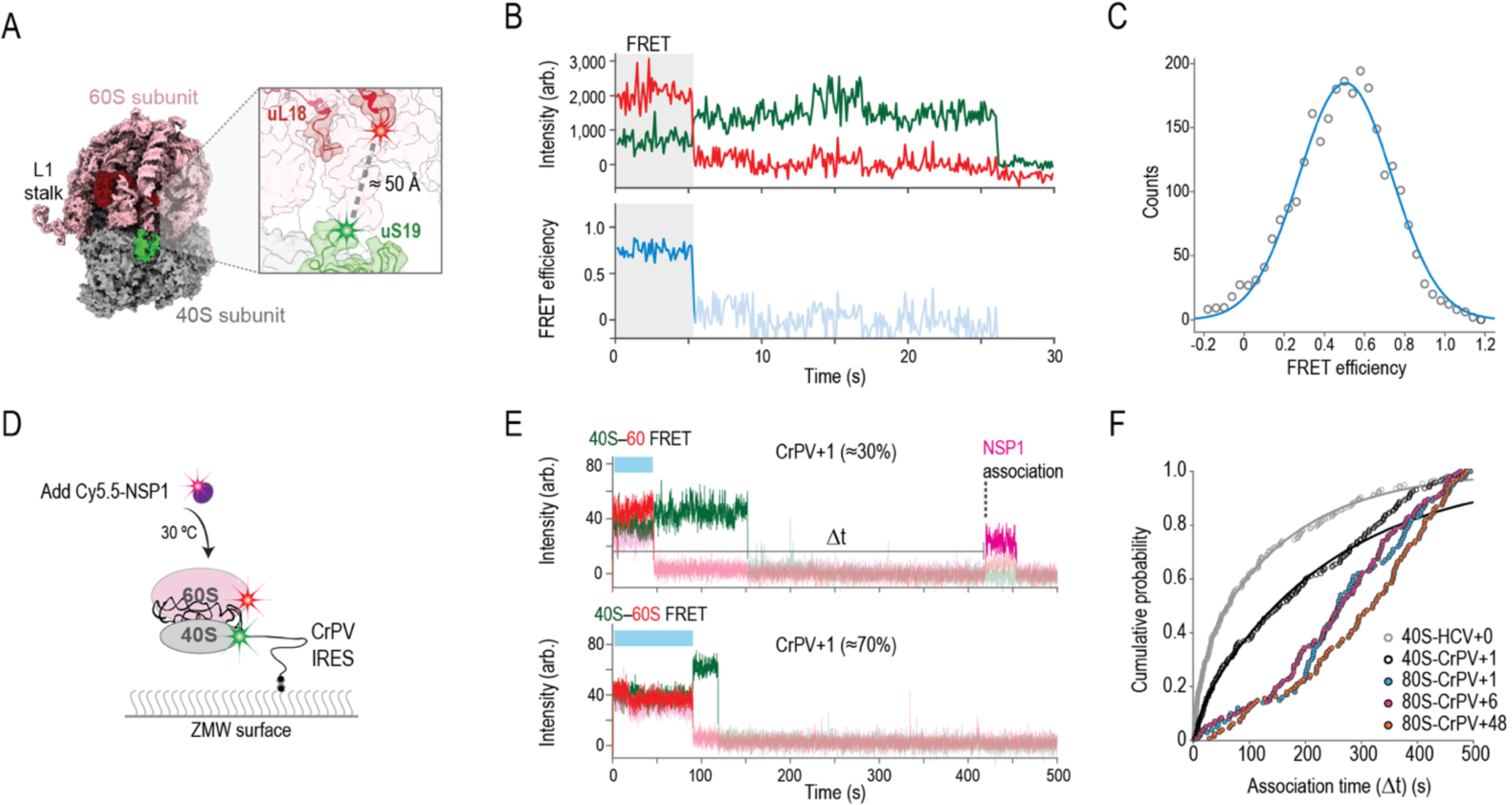
NSP1 inefficiently associated with 80S ribosomes assembled on the CrPV IRES. **A)**. Model of the human 80S ribosome (PDB: 4UG0). The 40S and 60S subunits are in gray and pink, respectively. The ybbR tag was fused to either the N-terminus of uS19 (green, 40S) or the C-terminus of uL18 (red, 60S). Based on available structural models, the ybbR tags were predicted to be within FRET distance in translation-competent 80S ribosomes. **B)**. Example fluorescent trace and calculated FRET efficiency plot. 40S-ybbR-Cy3 and 60S-ybbR-Cy5 subunits were incubated with the CrPV IRES to assemble 80S ribosomes on the biotinylated RNA. Following tethering of the complex, molecules were imaged at equilibrium using TIRFM. Molecules were expected to begin in a Cy3 (green, FRET donor) to Cy5 (red, acceptor) FRET state, followed by photobleaching of both dyes. The region of the trace that corresponds to FRET is highlighted by the gray box. **C)**. Plot of the distribution of observed FRET efficiencies for the inter-subunit FRET signal on 80S ribosomes. Frequencies of observed FRET efficiencies were binned into 35 bins (open circles) across the indicated range. The line represents a fit to a single-gaussian function, which yielded a mean FRET efficiency of 0.5 ± 0.01 (95% C.I.). *n = 104*. **D)**. Schematic of the single-molecule fluorescence experimental setup. 40S ribosomal subunits were labeled with Cy3 dye via uS19-ybbR, and 60S subunits were labeled with Cy5 via uL18-ybbR. Pre-formed 80S–CrPV IRES complexes were tethered to the ZMW imaging surface. At the start of data acquisition, Cy5.5-NSP1 (N-terminal ybbR tag) was added at 25 nM (final concentration) at 30 °C. **E)**. Example single-molecule fluorescence traces that depict addition of Cy5.5-NSP1 to tethered CrPV+1 RNAs bound by 80S ribosomes. The 40S subunit was labeled with Cy3 (green), 60S subunit with Cy5 (red), and NSP1 with Cy5.5 (magenta). The two traces are from the same experiment where 80S–CrPV+1 complexes were tethered. The top trace depicts a complex with an NSP1 binding event (∼30% of traces) and the bottom trace lacks an NSP1 event (∼70% of traces). Raw fluorescence intensities were corrected in this image to set baseline intensities to zero for presentation. Due to bleed through across the three fluorescent channels, the Cy3, Cy5, and Cy5.5 signals were made transparent before and after relevant events for presentation here. The association time (Δt) was defined as the time elapsed from the addition of Cy5.5-NSP1 until the burst of Cy5.5 fluorescence (magenta), which signified NSP1 association. The lifetime was defined as the duration of the Cy5.5 fluorescence signal. **F)**. Plot of the cumulative probability of observed NSP1 association times with the indicated ribosomal– CrPV IRES complexes at 30 °C. Cy5.5-NSP1 was added at 25 nM (final concentration). Lines represent fits to double-exponential functions. See **Supp. Table 4** for samples sizes and parameters for fits.

By leveraging the FRET signal and the CrPV IRES, we examined whether NSP1 associated with 80S ribosomes assembled on an mRNA (**Fig. 4D**). We generated RNAs as above with 1, 6, and 48 nucleotides downstream of the CCU codon present in the ribosomal A site (**Supp. Fig. 7A**). With these models, the RNA is shifted three nucleotides further into the entry channel relative to the HCV IRES^40^. Therefore, CrPV+6 and CrPV+48 will have mRNA that at least partially occludes the NSP1 binding site, while CrPV+1 will not. When added at 25 nM to 40S–

CrPV+1 complexes, we observed slower (median association time ≈ 119–189 s, 95% C.I.) and less efficient (45 ± 4%, 95% C.I.) Cy5.5-NSP1 association relative to that of 40S–HCV+0 complexes (median ≈ 46–82 s and 72 ± 4%, 95% C.I.) (**Supp. Fig. 7B,C** and **Supp. Table 4**). This finding likely reflects heterogeneity of the 40S subunit head conformation when bound to the CrPV IRES^41^, unlike the near-homogenous open conformation induced by the HCV IRES. Further inclusion of 60S subunits to yield 80S– CrPV+1 complexes inhibited NSP1 association (median association time ≈ 235–285 s, 95% C.I.), similar to the inhibition observed on both 80S–CrPV+6 and 80S–CrPV+48 complexes (median association times ≈ 243–292 s and 288–354 s, 95% C.I.) (**Fig. 4E,F** and **Supp. Fig. 7C**). Thus, even when mRNA was absent from it, the conformation of the mRNA entry channel on 80S–CrPV IRES complexes was incompatible with rapid NSP1 association.

Our findings support a model where NSP1 associates with the open conformation of the 40S ribosomal subunit, in the absence of mRNA within the entry channel. They also suggest that NSP1 is unable to rapidly bind 80S ribosomes in either the canonical or rotated states prior to peptide bond formation and translocation. Future studies will be needed to examine whether NSP1 accesses other states of the 80S ribosome.

### NSP1 remained bound to 40S subunits upon association with model mRNAs

Our data above indicated that mRNA inhibited NSP1 association with the 40S ribosomal subunit. However, it remained unclear whether RNA segments downstream of a start codon would destabilize the NSP1–40S subunit complex upon mRNA recruitment. To focus solely on NSP1, the 40S subunit, and mRNA, we again leveraged the HCV IRES. Using Cy5.5-NSP1 and 40S-Cy3 subunits, we pre-formed NSP1–40S complexes and added the complex at 15 nM to ZMWs with surface-immobilized HCV+0 or HCV+48 model mRNAs (**Supp. Fig. 8A-C**). On HCV+0 and HCV+48, NSP1 co-associated with 57 ± 6% and 60 ± 6% (99% C.I.) of 40S subunits (**Fig. 5A,B** and **Supp. Table 5**), which indicated near saturation of 40S subunits with NSP1. Association of the NSP1–40S subunit complex with the tethered RNAs had kinetics similar to 40S subunits alone (**Fig. 5C,D** and **Supp. Fig. 8D,E**). After association, NSP1 remained bound to the 40S subunit for approximately 60 s on average (median lifetimes 39–74 s and 52–82 s, 95% C.I.) (**Fig. 5D** and **Supp. Fig. 8F**). Consistent with our findings using the HCV IRES, NSP1 lifetimes on the ribosomal subunit were similar for both CrPV+1 and CrPV+48 model RNAs (**Supp. Fig. 8G-N**). Thus, the stability of NSP1 on the 40S subunit was unaffected upon direct association with the structured mRNAs we examined.

**Figure 5.**
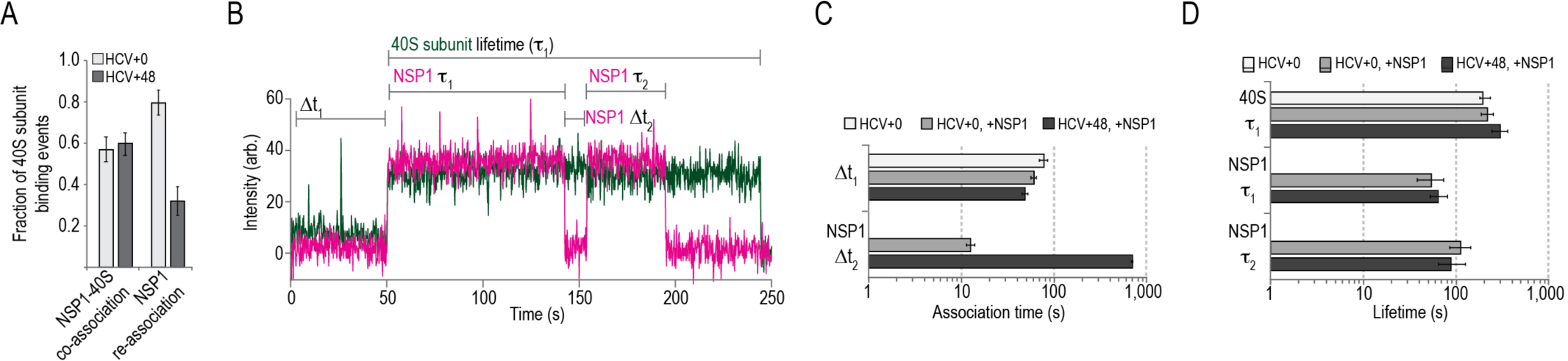
NSP1 remained bound to 40S subunits upon association with model mRNAs. **A)**. Plot of the fraction of 40S subunit binding events that had a co-associated NSP1 (left) or at least one NSP1 re-association event (right), as defined below in panel B. Error bars represent 99% C.I.. **B)**. Example single-molecule fluorescence trace that depicts association of NSP1–40S subunit complexes with a tethered HCV+0 IRES molecule. The 40S subunit and NSP1 were labeled with Cy3 (green) and Cy5.5 (magenta) dyes, respectively. Raw fluorescence intensities were corrected in this image to set baseline intensities to zero for presentation. The NSP1–40S subunit association time (Δt_1_) was defined as the time elapsed from the addition of the complex until the burst of Cy3 and Cy5.5 fluorescence, which signified association of the NSP1-40S subunit complex with the tethered IRES. In experiments that lacked NSP1, Δt_1_ was defined using the first burst of Cy3 signal alone. The 40S subunit lifetime (*τ*_1_) was defined as the duration of the Cy3 fluorescence signal. The initial NSP1 lifetime (NSP1 *τ*_1_) was defined as the duration of the Cy5.5 signal that co-appeared with the Cy3 signal. For NSP1 re-association analyses, we focused on ZMWs where a single 40S subunit associated within the first 200 s (≈ 75% of all events, see **Supp. Fig. 8D**). We then quantified the time elapsed from the loss of the first Cy5.5 signal to the next burst of Cy5.5 fluorescence at least ≈ 20 s in length (≈ 70% of initial NSP1 binding events, see **Supp. Fig. 8F**), which was defined as the NSP1 re-association time (NSP1 Δt_2_). The duration of this second Cy5.5 event was defined as the re-associated NSP1 lifetime (NSP1 *τ*_2_). **C**,**D)**. Plot of association times (panel C) and median lifetimes (panel D) from the indicated experiments. ‘+NSP1’ indicates experiments where pre-formed NSP1–40S subunit complexes were added. Association times (40S Δt_1_ and NSP1 Δt_2_) were defined as the inverse of the fast association rate derived from fits to double-exponential or linear functions, with error bars indicating the 95% C.I. of the rate. Error bars for median lifetimes (40S *τ*_1_, NSP1 *τ*_1_ and *τ*_2_) represent 95% C.I.. See **Supp. Table 5** for samples sizes and parameters for fits.

To delineate competition between NSP1 and mRNA for the 40S cleft, we next asked whether NSP1 could re-associate stably with single IRES–40S subunit complexes and how re-association was impacted by long segments of RNA downstream of the start codon. Indeed, following loss of the initial NSP1 signal (due to dye photobleaching or NSP1 departure), 80 ± 6% (99% C.I.) of 40S–HCV+0 complexes had at least one additional stable (≥ 20 s) NSP1 binding event (**Fig. 5A,B** and **Supp. Fig. 8B**). In contrast, only 32 ± 7% (99% C.I.) of 40S– HCV+48 complexes had a second, stable NSP1 event (**Fig. 5A,B** and **Supp. Fig. 8C)**. When multiple NSP1 association events were observed on a single 40S–IRES complex, the time that separated them was at least 55-fold longer on HCV+48 (apparent *k*_*on*_^*-1*^ ≈ ≥ 700 s) relative to HCV+0 (apparent *k*_*on*_^*-1*^ ≈ 13 ± 1 s, 95% C.I.) (**Fig. 5C** and **Supp. Fig. 8O**). The lifetimes of initial and re-associated NSP1 binding events were similar (**Fig. 5D** and **Supp. Fig. 8P**). Together, these findings indicated that once NSP1 dissociated from the 40S–HCV+48 complex, mRNA was accommodated more rapidly into the mRNA entry channel, thereby inhibiting re-association of NSP1.

## DISCUSSION

Shutdown of host protein synthesis is a common feature of viral infection. Most characterized mechanisms involve the covalent inactivation of key eIFs or their regulators (*e*.*g*., eIF2 and eIF4F^42^). Here, we provide insight into a distinct form of translation inhibition employed by SARS-CoV-2 and closely-related CoVs. The first protein encoded in the viral genomic RNA, NSP1, directly targets the small subunit of the human ribosome to inhibit protein synthesis. Based on our findings and recent structural studies^31,32^, we suggest that NSP1 preferentially associates with the open conformation of the 40S subunit to prevent proper accommodation of mRNA during translation initiation (**Fig. 6**).

**Figure 6.**
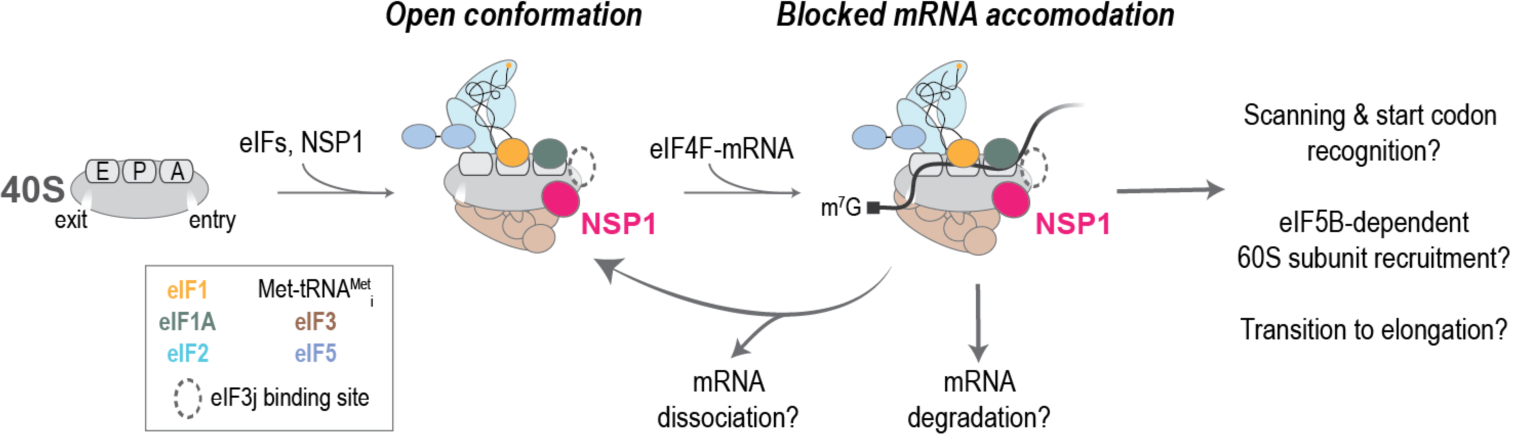
Proposed model. In the absence of eIF3j, NSP1 preferentially associates with the open conformation of the 40S subunit to block full accommodation of the mRNA in the entry channel, which inhibits translation initiation. How incomplete mRNA accommodation impacts mRNA recruitment, scanning, start codon selection, 60S subunit recruitment, and the transition to elongation remain open questions. Whether and how NSP1–40S subunit complexes lead to mRNA degradation is also unknown.

NSP1 is a potent inhibitor of human translation initiation. When we added purified NSP1 to human HeLa cell extract, we observed a dramatic reduction in translation of our model for human GAPDH mRNA. Inhibition was specific, as mutations in two NSP1 amino acids (KH164-165) necessary for 40S subunit binding abrogated the NSP1 inhibition. The apparent *IC*_*50*_ value for wild-type NSP1-mediated inhibition suggests near stochiometric association of NSP1 with 40S subunits in the cell extract, which agrees well with our best estimate for the *K*_*D*_ of the interaction (≤ 4 nM). Based on our kinetic analysis of protein synthesis, we demonstrate that such NSP1 association with the 40S subunit inhibits the initiation phase of translation. Consistently, a recent study^31^ illustrated that ectopic expression of SARS-CoV-2 NSP1 reduced the abundance of actively-translating polysomes and increased the abundance of 80S monosomes, a hallmark of translation initiation defects. Similar findings also have been observed with the SARS-CoV protein^16,17^.

NSP1 inhibited translation of SARS-CoV-2 model mRNAs at levels comparable to that for a model human mRNA. This finding suggests that NSP1 is a general inhibitor of protein synthesis. The high affinity of NSP1 for the 40S subunit likely demands buildup of NSP1 protein levels before translation is inhibited broadly, which may enable viral protein synthesis to proceed unimpeded during early stages of infection. Once NSP1 has accumulated, the increased translation efficiency of the viral mRNAs relative to human mRNAs we and others^32^ have observed may enable the virus to synthesize sufficient amounts of viral proteins, even when translation is largely shutdown. However, it is possible that another viral protein, a different segment of the viral genome, or another mechanism (*e*.*g*., sequestration) allows the virus to evade translation inhibition. Future studies in the context of infected human cells are needed to deconvolute these possibilities.

To inhibit translation initiation, multiple lines of evidence suggest that NSP1 preferentially associates with the open conformation of the 40S subunit. First, of all eIFs we examined, NSP1 association was enhanced the most (≈ 2-fold) by eIF1. This protein binds to the 40S subunit at the ribosomal P site^43–45^, where it has a critical role during start codon recognition^46–50^. Upon its association, eIF1 repositions the 40S subunit head, which concomitantly opens the mRNA entry channel^51,52^. Given that eIF1 and NSP1 binding sites are non-overlapping, our findings indicate that eIF1 allosterically enhances NSP1 association, likely by altering the conformation of the mRNA entry channel to allow NSP1 access. Second, the most rapid NSP1 association with the 40S subunit we observed was in the presence of the HCV IRES. This structured RNA directly manipulates the ribosomal subunit to bypass eIFs and initiate translation^53^. One of its flexible segments, domain II, makes extensive contacts with the 40S subunit head, which swivels, opening the mRNA entry channel^35–37^. In our assays, the estimated rate of NSP1 association with the HCV+0 model mRNA was 3–4 µM^-1^ s^-1^, nearly an order of magnitude faster than with 40S subunits alone. Enhanced NSP1 association with this complex relative to the eIF1–40S subunit complex likely reflects inherent differences in their stability. Whereas the IRES–40S subunit complex is quite stable (*K*_*D*_ ≈ 2 nM, *k*_*off*_ ≈ 0.002 s^-1^)^34,54^, the eIF1–40S subunit interaction is more closed conformations. Consistently, NSP1 associated more slowly with the 40S–CrPV+1 complex, which likely also contains a heterogenous mix of entry channel conformations^41^.

In striking contrast, NSP1 competes with mRNA to bind the ribosome. When 40S subunits were pre-incubated with an mRNA that had at least 12 nucleotides downstream of the start codon, we observed marked inhibition of NSP1 association with the ribosomal subunit. On such mRNAs, NSP1 contacts with helix 18 of the 18S rRNA and ribosomal proteins uS3 and uS5 in the mRNA entry channel are at least partially occluded by the accommodated mRNA. Consistently, when we pre-incubated mRNA in extracts prior to NSP1 addition, NSP1 failed to inhibit translation in our real-time protein synthesis assays. And in the reciprocal experiment, pre-incubation with NSP1 yielded a delay in protein synthesis (≈ 200 s) similar in length to our best estimate for the lifetime of the NSP1–40S subunit interaction (at least ≈ 240 s). In parallel, NSP1 association with the 40S subunit also was inhibited by pre-incubation with eIF3j, which binds near the mRNA entry channel^55,56^. The C-terminus of eIF3j may sterically block NSP1 association^55^ or eIF3j may limit movement of the 40S subunit head to promote a conformation of the entry channel inaccessible to NSP1, which may reflect the likely preference of eIF3j for the closed conformation of the 40S subunit^57^.

In further support of a competition model, NSP1 remained bound to the 40S ribosomal subunit upon recruitment of an mRNA— regardless of its length. This finding suggests that mRNA itself is insufficient to dislodge or destabilize the NSP1–40S subunit interaction. Based on our observed lifetime of that interaction (at least ≈ 240 s), NSP1 likely will remain associated with the ribosomal subunit for longer than the time frame of translation initiation on transient (*K*_*D*_ ≈ 50 nM, *k*_*off*_ ≈ 0.36 s^-1^)^28,50^, which many mRNAs (< 60 s)^58,59^, thereby blocking full would lead to a mixed population of open and accommodation of the mRNA. Whether NSP1 can be dislodged by other host proteins or eIFs, such as helicases eIF4A, DDX3X, or DHX29, remains unclear. Nevertheless, once NSP1 dissociated, mRNA was accommodated into the entry channel, which prevented re-association of NSP1. Thus, our findings suggest that the presence of NSP1 or mRNA within the mRNA entry channel of the 40S subunit are mutually exclusive. It is feasible that such competition blocks eIF4F-mediated recruitment of an m^7^G-capped mRNA to the 40S subunit or a subsequent step in initiation, which awaits elucidation by future studies. Collectively, our work provides a biophysical foundation for deeper understanding of NSP1-mediated shutdown of host translation and its impact on SARS-CoV-2 viral pathogenesis.

## ACKNOWLEDGEMENTS

We are grateful to Michael Lawson, Jan Carette, and other members of the Puglisi and Carette labs for helpful guidance, discussions, and feedback. We also appreciate discussions with and advice from Chris Fraser and Masa Sokabe. We thank Peter Sarnow and the Sarnow lab for sharing cell culture equipment, and the Stanford Shared FACS Facility for completion of all single-cell sorting. C.P.L. is a Damon Runyon Fellow supported by the Damon Runyon Cancer Research Foundation (DRG-#2321-18); A.G.J. was supported by a National Science Foundation Graduate Research Fellowship (DGE-114747); and J.W. was supported by a postdoctoral scholarship from the Knut and Alice Wallenberg Foundation (KAW 2015.0406). Research on eukaryotic translation in the laboratory of J.D.P. is funded by the National Institutes of Health (GM011378, AI047365, and AG064690).

## MATERIALS AND METHODS

### Molecular cloning

See **Supplementary Table 6** for all relevant sequences.

#### NSP1

Codon-optimized SARS-CoV-2 NSP1 and relevant mutants were cloned into a vector purchased from the UC Berkeley QB3 MacroLab (vector 1B) using their standard protocol. Synthetic DNA that encoded wild-type, RK124-125AA mutant, KH164-165AA mutant, and ybbR-tagged NSP1 sequences were purchased from Integrated DNA Technologies (IDT). All sequences were verified by Sanger sequencing. The resulting plasmids encoded NSP1 proteins tagged on the N-terminus with a 6-histidine tag followed by a TEV protease cleavage site (NH_2_-6His–TEV–NSP1-COOH). Wild-type NSP1 reference sequence (nts 266-805) was obtained from NCBI GenBank accession MN997409.1. When noted, a ybbR tag was included either on the N-terminus (ybbR-NSP1) or the C-terminus (NSP1-ybbR).

#### HCV IRES

A synthetic DNA was purchased from IDT that contained a 5’ flanking sequence (pUC19 backbone sequence), a T7 promoter (TAATACGACTCACTATAG), and the HCV IRES (nts 1-344, including the AUG codon). Downstream of the AU<u>G</u> included the rest of domain IV, the core sequence, and a 3’ extension: AGCAC<u>G</u>AATCC<u>T</u>AAACCTCAAAG<u>A</u>AAAACCGCCAGAACCATGGAAGA<u>C</u>. DNA templates that encoded HCV+0, +6, +12, +24, and +48 (relative to the ‘G’ of the start codon, 3’-end indicated by underlined nts) were generated via standard PCRs using NEB Phusion polymerase (25 cycles), a common 5’ primer (upstream of the T7 promoter), and specific 3’ primers.

#### CrPV IRES

A plasmid that encodes the IRES of the intergenic region of CrPV with the first codon (Ala) replaced with a Phe codon (TTC) was described previously^60^. The sequence from CCU (bolded) of the IRES to the 3’ end was: **CCT**<u>T</u>TCAC<u>A</u>TTTCAAGATACCGGCGCCATGGAAGACGCCAAAAACATAAA<u>G</u>. DNA templates were generated as for the HCV IRES, with the underlined nts representing the 3’-terminus. We selected +1 as the shortest segment to promote proper folding of the IRES.

#### GAPDH nanoLuciferase

The reporter was designed to contain a 5’ T7 promoter, the nLuc coding sequence (Promega) flanked by the 5’ and 3’UTRs from human GAPDH (NCBI GenBank accession: AF261085), a poly(A) tail, and PmeI and SpeI restriction enzyme consensus sites. The reporter construct was purchased from IDT as a plasmid with an pUCIDT backbone and propagated in the DH5*α* strain of *E. coli*. DNA sequence identity was confirmed by Sanger sequencing. The plasmid was linearized by restriction digest with SpeI (NEB, #R0133) for templated in vitro transcription.

#### SARS-CoV-2 nanoLuciferase

Viral DNA (NCBI GenBank accession MN997409.1) constructs were designed such that the nLuc coding sequence was flanked by either the full-length 5’UTR or the subgenomic 5’ leader sequence and one of two 3’’UTR sequences, and a poly A tail followed by a SpeI consensus sequence. The full-length 5’UTR incuded the first 27 nt of the viral ORF in order to maintain the predicted stem-loop structure within the 5’UTR. The 3’UTR (L) began after the N protein stop codon and the 3’UTR (S) began after ORF10. Synthetic DNAs were purchased as GeneBlocks from IDT (5’ leader constructs) or as Gene Parts from GenScript (5’UTR constructs). All synthetic DNAs had a 5’-terminal XmaI consensus sequence and 3’-terminal HindIII consensus sequence. Restriction digest cloning was used to insert viral reporter DNA into a pUC19 vector (NEB, #R10180S, #R3104S, #M2200S).

### NSP1 expression, purification, & labeling

NSP1 expression plasmids were transformed into OneShot BL21(DE3) cells (Invitrogen) and grown overnight at 37 °C on LB agar plates supplemented with 50 µg/mL kanamycin. Liquid cultures of single colonies were grown to OD_600_ ≈ 0.5 at 37 °C in LB supplemented with kanamycin. Cultures were shifted to 18 °C for 30 minutes, 0.5 mM IPTG was added, and cultures were grown for 16-20 h at 18 °C. Cells were harvested by centrifugation at 5,000 x *g* for 15 min at 4 °C in a Fiberlite F9 rotor (ThermoFisher, cat. # 13456093). Cells were lysed by sonication in lysis buffer (20 mM Tris-HCl pH 8.0, 300 mM NaCl, 10% (v/v) glycerol, 40 mM imidazole, and 5 mM β-mercaptoethanol), and lysates were cleared by centrifugation at 38,000 x *g* for 30 min at 4 °C in a Fiberlite F21 rotor followed by filtration through a 0.22 µm syringe filter. Clarified lysate was loaded onto a Ni-NTA gravity flow column equilibrated in lysis buffer, washed with 20 column volumes (CV) of lysis buffer, 20 CV of wash buffer (20 mM Tris-HCl pH 8.0, 1 M NaCl, 10% (v/v) glycerol, 40 mM imidazole, and 5 mM β-mercaptoethanol), and 10 CV of lysis buffer. Recombinant proteins were eluted with five sequential CV of elution buffer (20 mM Tris-HCl pH 8.0, 300 mM NaCl, 10% (v/v) glycerol, 300 mM imidazole, and 5 mM β-mercaptoethanol). Fractions with recombinant protein were identified by SDS-PAGE analysis. The relevant fractions were dialyzed overnight at 4 °C into ybbR-labeling buffer (50 mM HEPES-KOH pH 7.5, 250 mM NaCl, 10 mM MgCl_2_, 10% (v/v) glycerol, and 1 mM DTT) or TEV Buffer (20 mM Tris-HCl pH 8.0, 250 mM NaCl, 10% (v/v) glycerol, 10 mM imidazole, and 5 mM β-mercaptoethanol), as appropriate. Fluorescent labeling via the ybbR tag was performed essentially as described^26,29^. Briefly, 14.5 µM ybbR-NSP1 or NSP1-ybbR were incubated at 37 °C for 90 min with 4 µM Sfp synthase enzyme and 20 µM of either Cy3-CoA, Cy5-CoA, or Cy5.5-CoA substrates, respectively. Free dye was removed via purification over 10DG-desalting columns (Bio-Rad, cat.# 7322010) equilibrated in TEV buffer. Dye-labeled and non-labeled NSP1 proteins were incubated with TEV protease at 22 °C for 1.5 hrs followed by 30 min at 37 °C. TEV protease, Sfp synthase, and the cleaved 6His tag were removed via a subtractive Ni-NTA gravity column equilibrated in TEV buffer, with the flow-through collected. NSP1 proteins were subjected to a final purification step using size exclusion chromatography (SEC) on a Superdex 75 column (23 mL) equilibrated in SEC buffer (20 mM HEPES-KOH pH 7.5, 250 mM KOAc, 10% (v/v) glycerol, and 1 mM DTT). Fractions containing NSP1 were concentrated using a 10 kD MWCO Amicon Ultra centrifugal filter, aliquoted, flash frozen on liquid N_2_, and stored at −80 °C. Protein concentrations were determined via absorption at 280 nm using a nanodrop for total protein, and at 548 nM or 646 nM for Cy3 or Cy5/5.5 labeled proteins, respectively. For ybbR-NSP1, labeling efficiencies were 50-70%. For NSP1-ybbR, the labeling efficiency was much lower (<20%), and the protein had reduced translation inhibition activity, which is why it was excluded from single-molecule analyses.

### nLuc *in vitro* translation assays

HeLa cell-free translation (ThermoFisher, #88884) reactions setup according to manufacturer’s protocol were programed with a final mRNA concentration of 200 nM (endpoint) or 80 nM (real-time). For reactions containing NSP1, an equal volume of NSP1 buffer was added to the paired no NSP1 control reaction. Prism8 was used for dose-response analysis ([inhibitor] vs. response with variable slope (four parameters) nonlinear fit) and statistical analysis (one-way ANVA Turkey’s multiple comparisons test, unpaired t-test).

#### Endpoint assays

IVT reactions were incubated at 37°C for 45 min and then immediately transferred to an ice water bath and diluted 1:1 with cold Glo Lysis Buffer (Promega, #E2661). All samples were brought to room temperature and mixed with a 1:1 volume of nGlow solution (Promega, *#*N1110). Samples (90% of total volume) were loaded into non-adjacent wells of a 384-well plate. Sample luminescence was measured 7 min post nGlow solution addition using a BioTek Neo2 multi-mode plate reader (25°C, 114LUM1537 filter, gain of 135). Luminescence signal was monitored for an additional 30 min at 3 min intervals to verify luminesce signal of all samples decayed at the same rate.

#### Real-time assays

HeLa IVT reactions were prepared by addition nGlow substrate to the cell-free translation mix with a 1:10 v/v ratio. Before the addition of mRNA and/or NSP1, the IVT reactions were transferred to non-adjacent wells in a 384-well plate and equilibrated to 30°C in the plate reader. All other reagents were maintained at 30°C and then added to the IVT reactions according to the order-of-addition assay schematic outlined in **Fig. 1D**. The preincubation (30°C, 2 min) was performed in the plate reader. Kinetic monitoring of the samples (36 min, 15 s intervals) was initiated during the equilibration step. Reagent additions and plate transfer times were noted during the experiment and used to post-synchronize *t*_0_ to mRNA addition. Data was analyzed in MatLab using the approach developed by Vassilenko *et al*.^25^. The raw data was smoothed using Savitzky-Golay filtering (frame length ≤ 5, 2^nd^ degree polynomial) to maintain the shape and magnitude of the raw data. The plateau value of the 1^st^ derivative of the smoothed curve is equal to the synthesis rate^25,61^. Mean synthesis time, which is lag time between mRNA addition and the initial detection of nLuc signal, is the sum of initiation, elongation, and termination time. Synthesis times were extracted from the data by fitting the 2^nd^ derivative of the smoothed data to a Gaussian distribution using the equation

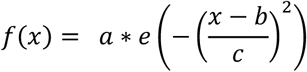

where *a* = amplitude, *b* = centroid (median), *c* = variance^25,62^. Statistical analysis of the real-time nLuc activity was done as described above.

### Native gel assays

Native composite agarose-acrylamide gels were prepared as described^63^. Briefly, 2.75% acrylamide (37.5:1), 0.5% Nusieve GTG agarose composite gels were prepared in the following buffer: 25 mM Tris-OAc pH 7.5, 4 mM KOAc, 2 mM Mg(OAc)_2_, 0.5 mM DTT, 2.5% glycerol, 0.1% (v/v) TEMED, and 0.1% (v/v) fresh ammonium persulfate. The gels were cooled at 4 °C for 20 minutes and allowed to further polymerize at room temperature for 90 min. Prior to removal of combs, gels were placed at 4 °C for 15 min. Gels were run in ice-cold running buffer (25 mM Tris-OAc pH 7.5, 4 mM KOAc, and 2 mM Mg(OAC)_2_) for 30-60 min at 4 °C. All assays were repeated at least three times. For complex formation, the indicated components were incubated in ribosome assay buffer (30 mM HEPES-KOH pH 7.4, 100 mM KOAc, 2 mM Mg(OAc)_2_) at 37 °C for 15 minutes. Unless noted, NSP1 with a C-terminal ybbR tag conjugated to a Cy5 dye was used in all gel shift experiments, as C-terminally tagged NSP1 from SARS-CoV was functional in cellular assays^12^ and Cy5 provides cleaner signal upon image acquisition with a Typhoon imager. For competition experiments, the competitor protein was pre-incubated with ribosomal subunits at 37 °C for 15 minutes, prior to addition of the labeled protein.

### Purification of human eIFs

#### eIF1 and eIF1A

Plasmids for the expression of recombinant human eIF1 and eIF1A were gifted by Christopher S. Fraser (UC Davis). Detailed cloning and recombinant protein expression and purification methods have been described previously^55^. Changes made to the eIF1 and eIF1A purification scheme include use of a HiTrap SP-HP (Cytiva Lifesciences, #17115201) column for the IEX chromatography step and the addition of a SEC step in which fractions eluted from the IEX column containing either eIF1 or eIF1A, as determined by SDS-PAGE analysis, were pooled and passed over a Superdex 75 10/300 (Cytiva, #17-5174-01) column to remove protein oligomers/aggregates and exchange purified factors into a storage buffer (20 mM Hepes, pH 7.5, 250 mM KOAc, 1 mM dithiothreitol (DTT), and 10% glycerol (v/v)). Purified factors were concentrated to ∼ 200 μM using a Millipore centrifugal filter and stored at −80°C.

#### eIF2 and eIF3

Endogenous initiation factor complexes were purified from 400 mL of HeLa post-nuclear extract that was kindly donated to our group by Robert Tijan (UC Berkeley) using previously detailed methods^55,64^. Additional guidance was received from Christopher S. Fraser (UC Davis).

#### eIF5

The expression and purification of recombinant eIF5 was carried out using a using a modified protocol based on previously established methods^65^. Alterations to the protocol included substitution of the MonoQ column with a HiTrap Q HP column (Cytiva Lifesciences, #29051325) for the anion exchange chromatography step, and adding a SEC step using a Superdex 200 10/300 (Cytiva Lifesciences, 28-9909-44) column to exchange purified eIF5 in to a storage buffer containing 20 mM Hepes, pH 7.5, 250 mM KOAc, 1 mM DTT, and 10% glycerol (v/v).

#### eIF3j

Human eIF3j was expressed and purified as done for NSP1, with the following changes. Human eIF3j was codon optimized for expression in *E. coli* and cloned with an N-terminal 6xHis tag and TEV protease cleavage site into a pET28b vector at the NcoI site. Protein was expressed by induction with 0.5 mM IPTG overnight at 17 °C. Following cleavage of the 6xHis tag and the subtractive Ni-NTA step, eIF3j was subjected to a final purification step using SEC on a 23 mL Superdex 75 10/300 (Cytiva, #17-5174-01) column equilibrated in storage buffer (20 mM HEPES-KOH pH 7.5, 150 mM KOAc, 10% (v/v) glycerol, and 1 mM DTT). Fractions with purified eIF3j were identified via SDS-PAGE, concentrated, flash frozen, and stored at −80 °C.

#### tRNA_i_

Human tRNA_i_ was transcribed from a DNA template with a 5’-end T7 promoter and hammer head ribozyme^55^. The plasmid was linearized via digestion with BstNI for transcriptional termination at the 3’-CCA of tRNA_i_. Purified and linearized plasmid was used as the template for in vitro transcription with T7 polymerase at 37 °C for 4 hrs at 16 mM MgCl_2_, during which the ribozyme self-cleaved (>80% efficiency). Mature tRNA_i_ was separated from pre-cursor RNA and cleaved ribozyme via 10% acrylamide gel electrophoresis in the presence of 8M urea. After excision of the tRNA_i_ band, the RNA was extracted 3x at room temperature for 12 hrs with 300 M ammonium acetate, and ethanol precipitated. tRNA_i_ was resuspended in 10 mM NaCl,10 mM Bis-tris, pH 7.0 and stored at −80°C. To charge tRNA_i_ with methionine, 100 µL of 60μM human tRNAi transcript was mixed with 618.6 µL ddH2O, and heated at 95 °C for 2 min, then immediately chilled on ice for 5 min. The resulting solution was mixed with 200 µL of 5x charging buffer (200 mM Tris-HCl pH7.5, 50 mM Mg(OAc)_2_, 5 mM DTT), 20 µL 100 mM ATP, 30 µL 10 mM L-methionine, 2 µL 1 M Mg(OAc)_2_ and 29.4 µL 34 µM yeast MetRS^66^, and incubated at 30 °C for 45 min. The charging reaction was stopped by addition of 100 µL 3M NaOAc pH5.2. The resulting tRNA was purified by phenol-chloroform-isoamyl alcohol (25:24:1, pH 5.2) extraction and ethanol precipitation. The pellet was resuspended with tRNA storage buffer (10 mM NaOAc pH 5.2, 50 mM Mg(OAc)_2_) and further purified by passing through BioRad P-6 columns that were equilibrated with tRNA storage buffer. The charging efficiency was ≈70% based on acid urea PAGE analyses^67^ of the final tRNA product.

### Real-time single-molecule assays using ZMWs

#### ZMW-based imaging

All real-time imaging was conducted using a modified Pacific Biosciences RSII DNA sequencer, which was described previously^30^. Unless otherwise noted, Cy3 dyes were excited using the 532 nm excitation laser at 0.6 µW/µm^2^. Cy5 and Cy5.5 dyes were excited with the 642 nm laser at 0.1 µW/µm^2^. In nearly all experiments, data were collected at 10 frames per second. The exception was when the 532 nm laser was used at 0.16 µW/µm^2^; in this case, data were collected at 3 fps to increase signal to noise ratios. ZMW chips were purchased from Pacific Biosciences. Prior to imaging, all ZMW chips were washed with 0.2% Tween-20 and TP50 buffer (50 mM Tris-OAc pH 7.5, 100 mM KCl). Washed chips were coated with neutravidin by a 5 min incubation with 1 µM neutravidin diluted in TP50 buffer supplemented with 1.3 µM of DNA blocking oligos and 0.7 mg mL^-1^ UltraPure BSA. Chips were washed with TP50 and ribosome assay buffer (30 mM HEPES-KOH pH 7.4, 100 mM KOAc, 2 mM Mg(OAc)_2_). For imaging, the ribosome assay buffer was supplemented with casein (62.5 µg mL^-1^) and an oxygen scavenging system^68^: 2 mM TSY, 2 mM protocatechuic acid (PCA), and 0.06 U/µL protocatechuate-3,4-dioxygenase (PCD). In all real-time single-molecule experiments, fluorescently-labeled NSP1 with an N-terminal ybbR tag was used, since it had translation inhibition and 40S-binding activities similar to the wild-type protein.

#### Tethered eIF1–40S and eIF1–eIF3j–40S complexes (Figure 2)

Prior to tethering, 200 nM biotin-40S subunits were incubated in ribosome assay buffer with 6 µM eIF1 at 37 °C for 20 min. For tethering, 1 nM of pre-formed complexes (by biotin-40S) were incubated on the neutravidin-coated ZMW surface for 15 min at room temperature. Non-tethered complexes were washed away with ribosome assay buffer prior to imaging. Upon start of data acquisition and in the presence of 1 µM eIF1, Cy3-NSP1 was added at the indicated concentrations and temperatures. In experiments with eIF3j, 2.5 µM eIF3j was pre-incubated for ∼5 min with tethered ribosomal complexes prior to addition of ybbR-NSP1 labeled with Cy3 dye (‘Cy3-NSP1’). Temperatures and laser powers were as indicated.

#### Tethered ribosomal pre-initiation complexes (Figure 2)

Complexes were formed by incubating 40 nM biotin-40S subunits in ribosome assay buffer with the indicated eIFs at 37 °C for 15 min. eIF concentrations were: 1 µM for eIF1, eIF1A, & eIF5; and 200 nM for TC-GMPPNP & eIF3. Tethering was conducted as above. As indicated, imaging was conducted in the presence of: 1 µM eIF1, eIF1A, & eIF5; 100 nM TC-GMPPNP; and 50 nM eIF3. Upon start of data acquisition at 30 °C, Cy3-NSP1 was added at 25 nM final concentration.

#### Tethered 40S–HCV IRES complexes (Figure 3)

The indicated HCV IRES RNAs were diluted to 200 nM in refolding buffer (20 mM cacodylate-NaOH pH 7.0, 100 mM KCl, and 1 mM EDTA), heated to 95 °C for 2 min, and slow cooled to room temperature (∼45 min). Once cooled, 4 mM Mg(OAc)_2_ was added to quench the EDTA, and RNAs were stored on ice until use. To form complexes, 75 nM of 40S-RACK1-Cy5 subunits were incubated in ribosome assay buffer with 20 nM of the indicated IRES RNAs at 37 °C for 20 min. As indicated (‘+eIFs’), 1 µM eIF1, 1 µM eIF1A, 1 µM eIF5, 300 nM TC-GMPPNP, and 240 nM eIF3 were included. Tethering was performed as above, except 1 nM of complex was incubated for 5 min at room temperature. Upon start of data acquisition at 30 °C, Cy3-NSP1 was added at 25 nM final concentration. As indicated (+eIFs), data acquisition was performed in the presence of 1 µM eIF1, 1 µM eIF1A, 1 µM eIF5, 200 nM TC-GMPPNP, and 150 nM eIF3.

#### Tethered M+41–40S subunit complexes (Figure 3)

The biotinylated model mRNA (M+41) and Cy5-labeled model mRNA (M+41-Cy5) were described previously^38^. To form complexes with M+41, 40 nM 40S-RACK1-Cy5 subunits were incubated in ribosome assay buffer with 500 nM M+41, 1 µM eIF1, 1 µM eIF1A, 1µM eIF5, and 400 nM TC-GMPPNP at 37 °C for 20 min. Tethering was performed as above, except 1 nM of complex was incubated for 10 min at room temperature. Upon start of data acquisition at 30 °C, Cy3-NSP1 was added at 25 nM final concentration in the presence of 1 µM eIF1, 1 µM eIF1A, 1 µM eIF5, and 200 nM TC-GMPPNP. For M+41-Cy5, conditions were identical except TC-GMPPNP was present at 200 nM and 100 nM during complex formation and imaging, respectively; and, eIF3 was included at 200 nM and 50 nM. eIF3 was included in these experiments to promote complex formation, which was inefficient.

#### Tethered 40S–CrPV and 80S–CrPV complexes (Figure 4)

CrPV IRES RNAs were refolded as with the HCV IRES. To form 40S–CrPV IRES complexes, 40S-uS19-Cy3 subunits at 75 nM were incubated in ribosome assay buffer with 20 nM of the indicated RNA at 37 °C for 15 min. To form 80S–CrPV IRES complexes, 75 nM 40S-uS19-Cy3, 150 nM 60S-uL18-Cy5, and 20 nM RNA were incubated. Tethering was performed as with the 40S– HCV complexes. Upon start of data acquisition at 30 °C, Cy5.5-NSP1 was added at 25 nM final concentration.

#### Addition of NSP1–40S subunit complexes (Figure 5)

HCV and CrPV IRES RNAs were refolded as above. To form NSP1–40S subunit complexes, 550 nM Cy5.5-NSP1 was incubated in ribosome assay buffer with 275 nM 40S-uS19-Cy3 subunits at 37 °C for 15 min. To tether biotinylated RNAs, 0.16 nM of refolded IRES was incubated on the neutravidin-coated ZMW surface for 5 min. Upon start of data acquisition at 30 °C, either 15 nM of 40S-Cy3 subunits alone or 15 nM of Cy5.5-NSP1–40S-Cy3 pre-formed complexes (final concentrations) were added.

#### Data analysis

Experimental movies that captured fluorescence intensities over time were processed using custom MATLAB scripts, as described previously^29,30^. Briefly, ZMWs with the desired fluorescence signals were identified by filtering for the signals at appropriate time points. Individual binding events were assigned manually based on the appearance and disappearance of the respective fluorescence signals. Unless noted, only the first NSP1 binding event longer than 4-5 s that occurred within the first 500 s of imaging was analyzed. Unless intractable due to complex stability or inhibition, approximately 200-300 single molecules were analyzed, indicated by single-step photobleach events. Association times were defined as the time elapsed from the addition of the labeled component until a burst of fluorescence for that component. The time of addition is controlled by the instrument and varies from experiment to experiment, but typically occurs within the first 10 s of data acquisition. Lifetimes were defined as the duration of the corresponding fluorescence signal.

Kinetic parameters were extracted by fitting observed data to single- or double-exponential functions as described^30^. On some complexes, the presence of a large, slow association phase made it difficult to derive reliable rates, as amplitudes for the association rates are assigned semi-arbitrarily during the fits. When this occurred, comparisons of median association times were used instead, which better reflected the raw data. In the results section, this is indicated by ‘median association times’, which only pertains to the indicated final concentration of NSP1. When possible, observed association rates derived from fits to exponential functions were reported, which are indicated by ‘apparent *k*_*on*_^*-1*^’ and were converted to times to facilitate comparisons across experiments where medians are reported. Regardless, all derived association rates, median association times, lifetimes, and the number of molecules examined are reported in **Supplementary Tables 2-5**. As indicated in the tables, fits to linear functions were used to estimate very slow association rates observed when NSP1 association was inhibited.

#### Statistical analyses

To calculate errors for NSP1 binding efficiency (*e*.*g*., Figure 3E), bootstrap analyses (n = 10,000) were performed to calculate 99% confidence intervals (C.I.) for the observed proportions using R. To calculate errors for median association times and lifetimes, bootstrap analyses (n = 10,000) were performed to calculate 95% C.I. of the observed median using MATLAB. Reported errors for derived rates represent 95% C.I. yielded from fits to linear, single-exponential, or double-exponential functions, as indicated.

### Equilibrium single-molecule analyses using total internal reflection fluorescence microscopy (TIRFM)

Our home-built, prism-based TIRFM instrument has been described previously^68–70^. All data were collected at room temperature at 10 frames per second with the EM gain set to 650. To form NSP1–40S subunit complexes, 40 nM biotin-40S subunits were incubated in ribosome assay buffer with 500 nM NSP1-ybbR-Cy5 and 6 µM eIF1 at 37 °C for 15 min. Pre-formed complexes were tethered to the surface of a neutravidin-coated quartz slide as with ZMWs above. Buffer conditions during TIRFM imaging were identical to those with ZMWs. During imaging via excitation with the 647 nm laser, 1 µM eIF1 and 4 nM NSP1-Cy5 (C-terminal ybbR tag) were present. To form 80S–IRES complexes, 100 nM of refolded CrPV IRES^60^ biotinylated on the 3’-terminus was incubated in ribosome assay buffer with 250 nM 40S-uS19-Cy3 and 750 nM 60S-uL18-Cy5 subunits. The complex was tethered as above at 1 nM. Emission data were collected in both the Cy3 (donor) and Cy5 (acceptor) channels following excitation of the Cy3 donor dye with the 532 nm laser. Co-localized molecules were identified and analyzed using custom MATLAB scripts. FRET events were assigned manually to a single state, with single molecules indicated by single-step photobleach events of the donor fluorophore. Background fluorescence intensities were corrected and normalized, and the gamma-corrected FRET efficiency distribution was calculated as described^71^.

### Ribosome purifications and labeling

40S and 60S ribosomal subunits were purified from the indicated cell lines and labeled with biotin or dyes as described^29^. To specifically install biotin on the ribosomal subunit, purified 40S-RACK1-ybbR subunits were incubated with 4-fold molar excess PEG_11_-biotin (ThermoFisher #21911) conjugated to co-enzyme A (CoA) and 2-fold molar excess Sfp synthase enzyme. To install fluorophores, purified 40S-RACK1-ybbR, 40S-uS19-ybbR, and 60S-uL18-ybbR subunits were incubated with 4-fold molar excess Cy3 or Cy5 dyes conjugated to CoA and 2-fold molar excess of Sfp synthase. All reactions proceeded for 90 min at 37 °C. To remove free biotin, dyes, and/or Sfp synthase, reactions were layered onto a low-salt sucrose cushion (30 mM HEPES-KOH pH 7.5, 100 mM KOAc, 0.5 M sucrose, 5 mM Mg(OAc)_2_, and 2 mM DTT), and centrifuged at 287,582 x *g* (90,000 rpm) for 1 hr at 4 °C in a TLA100.2 rotor (Beckman, ref. 362046) in 11×34 mm thick-wall polycarbonate ultracentrifuge tubes (Beckman, ref. 343788). Ribosome pellets were washed once and subsequently resuspended with ribosome storage buffer (30 mM HEPES-KOH pH 7.4, 100 mM KOAc, 5 mM Mg(OAc)_2_, 6% (v/v) sucrose, and 2 mM DTT), aliquoted, flash frozen in liquid N_2_, and stored at −80 °C.

### RNA in vitro transcriptions

#### HCV & CrPV IRES RNAs

Purified PCR products were used as the templates for in vitro transcription with the T7 MEGAScript T7 Transcription kit (ThermoFisher #AMB13345). Standard reaction conditions were used, except each nucleotide was present at a final concentration of 4 mM and the reaction was supplemented with 6 mM of 5’-biotin-G-Monophosphate (Trilink, #N-6003). Reactions were incubated at 37 °C for 2 hours. Transcribed RNAs were purified using the GeneJET RNA Purification Kit (ThermoFisher, #K0732).

#### Reporter mRNAs for IVT assays

Linearized plasmids were used as templates for in vitro transcription with the MessageMAX T7 ARCA-Capped Message Transcription Kit (Cell Script, # CMMA60710). Transcription reactions were setup using standard conditions and then incubated at 37°C for 30 min. Transcripts were purified using the MEGAclear Transcription Clean-up Kit (ThermoFisher, AM1908). The integrity and homogeneity of the reporter mRNAs were verified via native- and denaturing-PAGE.

### CRISPR-Cas9 & Homology-directed repair

#### Sequences

To generate the 40S-uS19-ybbR and 60S-uL18-ybbR cell lines, the guide sequences CGCAACACTCACCATCTTGC (uS19/RPS15) and AAAAATCATAGAAAATTGCT (uL18/RPL5) were cloned into the pX458 vector backbone using the published approach^72^. To insert the tandem ybbR and flag tags onto the endogenous copies of the genes, single-stranded DNA ultramer repair templates were purchased from Integrated DNA Technologies that contained about 40-60 nts of flanking sequence on either side of the desired insertion. See **Supplementary Table 6** for all guide oligo, repair template, and PCR screening oligo sequences.

#### Transfections, sorting, & screening overview

Approximately 24 h post seeding in a well of a 6-well plate, low-confluency (∼30%) wild-type HEK293T cells were transiently transfected (Liopfectamine 3000, ThermoFisher) with 1 µg of the relevant pX458 plasmid and 2 µg of ssDNA repair template. Cells were allowed to recover for 48 hrs. Single, eGFP-positive cells were sorted at the Stanford Shared FACS Facility into a well of a 96-well plate that contained 50% conditioned DMEM(high glucose) medium. Individual colonies were allowed to recover until they were visible by eye (approximately 1.5-2 weeks), upon which colonies were transferred to a well of a 24-well plate. Once confluent, colonies were screened via PCR, Sanger sequencing, and Western blot analyses. Successfully edited cell lines were expanded and ultimately stored as stocks in 5% DMSO-FBS solution.

#### PCR screening & Sanger sequencing

Genomic DNA was isolated from candidate cell lines using the QuickExtract DNA Extraction Solution (Lucigen, #QE09050) essentially as described, except 10 µL of resuspended cells were added to 50 µL of QuickExtract solution. Following extraction, 1 µL of gDNA was added to a standard 24 µL 2X GoTaq Green PCR (Promega, #M7123), typically ∼35 cycles. Sequences for uS19 and uL18 screening oligos are available in **Supplementary Table 6**. Following amplification, PCRs were analyzed using 2% agarose gel electrophoresis. PCR products with the desired insertion size were submitted for Sanger sequencing, following their purification.

#### Western blots

Whole cell lysates were analyzed via 4-20% SDS-PAGE followed by transfer to PVDF membranes. All antibodies were diluted in 5% Blotting-Grade Blocker (BioRad, #1706404) in TBST buffer. The following primary antibodies were used: anti-RPS15/uS19 at 1:500 from Abcam (ab168361); and, anti-RPL5 at 1:500 from GeneTex (GTX101821). Incubations with primary antibodies were for > 12 h at 4 °C. The secondary antibody was GeneTex Rabbit IgG antibody (HRP) (GTX213110-01) and was used at 1:10,000. Blots were imaged via chemiluminescence.

### Polysome profiling assays

Polysome profiling assays were performed as described^29^ and in duplicate for the indicated cell lines.

### Cell lines and growth

HAP1 cells that express RACK1-yybR were described previously^29^. HAP1 cells were grown at 37 °C with 5% CO_2_ in Iscove’s Modified Dulbecco’s medium (IMDM) (Gibco, #sh30228) supplemented with 10% (v/v) fetal bovine serum, 2 mM L-glutamine (Gibco), and 1X penicillin/streptomycin (Gibco). Wild-type HEK293T cells were purchased from ATCC (CRL-3216) and engineered HEK293T cells are described here. All HEK293T cell lines were grown as HAP1 cells except Dulbecco’s Modified Eagle’s Medium (DMEM) High Glucose medium (Gibco, #11965092) was used instead. Cell lines were not tested for mycobacterium.

### Data sharing plan

All plasmids, data, and custom analysis scripts are available upon reasonable request.

### Structural models

All structural models were rendered using ChimeraX^73^. The following PDB models were used: 4UG0^74^, 5A2Q^35^, and 6ZLW^31^.

## SUPPLEMENTARY FIGURES

**Supp. Figure 1.**
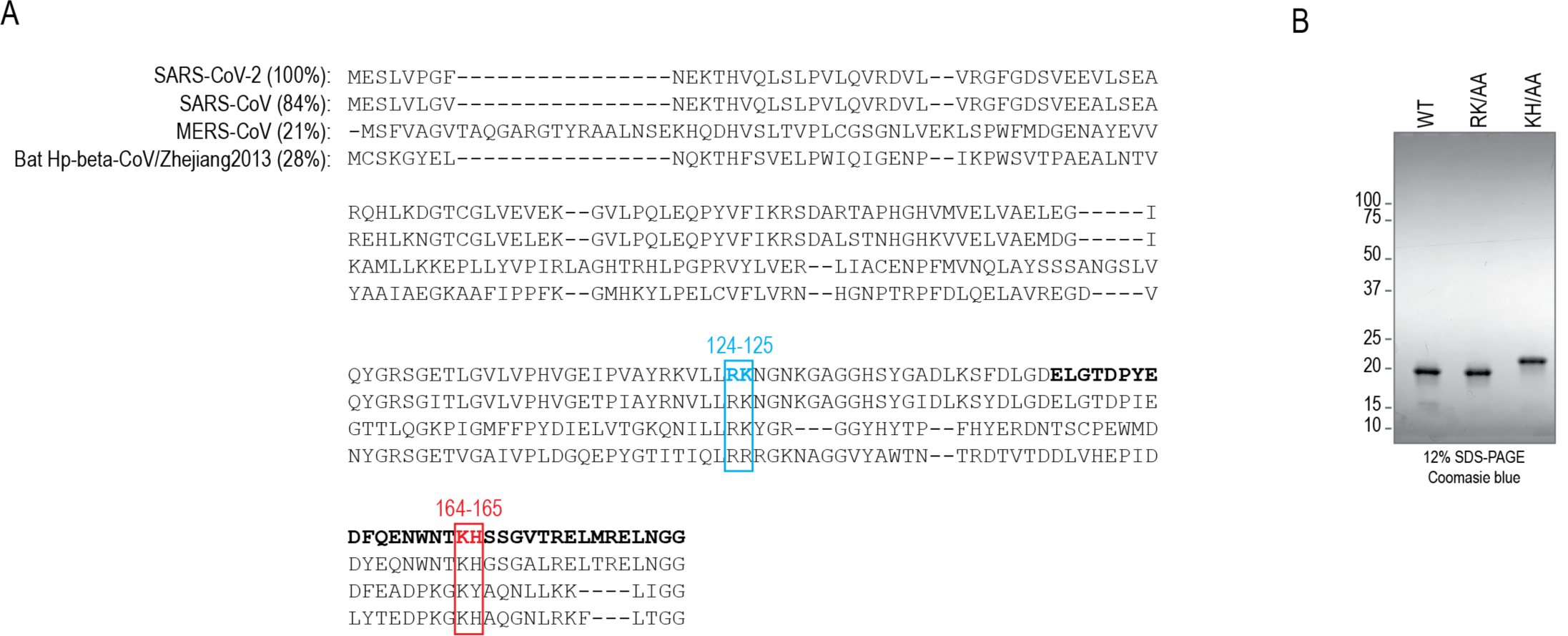
Conservation of NSP1 from CoVs. **A)**. Alignment of NSP1 protein sequences from SARS-CoV-2, SARS-CoV, MERS-CoV, and a related bat beta-CoV from the Hibecovirus subgenus (Bat Hp-beta-CoV/Zhejiang2013). The percent sequence identities are indicated (relative to SARS-CoV-2). The location of substitutions predicted to disrupt NSP1-mediated destabilization of mRNAs (RK124-125, blue box) and 40S-binding activities (KH164-164, red box) are highlighted. Based on recent structural studies, the C-terminal amino acids that dock within the mRNA entry channel of the 40S subunit are bolded. NCBI GenBank accessions are, respectively: MN997409.1, AY508724, JX869059, and KF636752. **B)**. Image of a gel that depicts recombinantly-expressed and purified SARS-CoV-2 NSP1 proteins examined using SDS-PAGE on a 12% acrylamide gel. The NSP1 mutant protein with KH164-165 substituted to alanine (KH/AA) runs at a different apparent molecular weight relative to the wild-type (WT) or RK124-125 to alanine (RK/AA) mutant.

**Supp. Figure 2.**
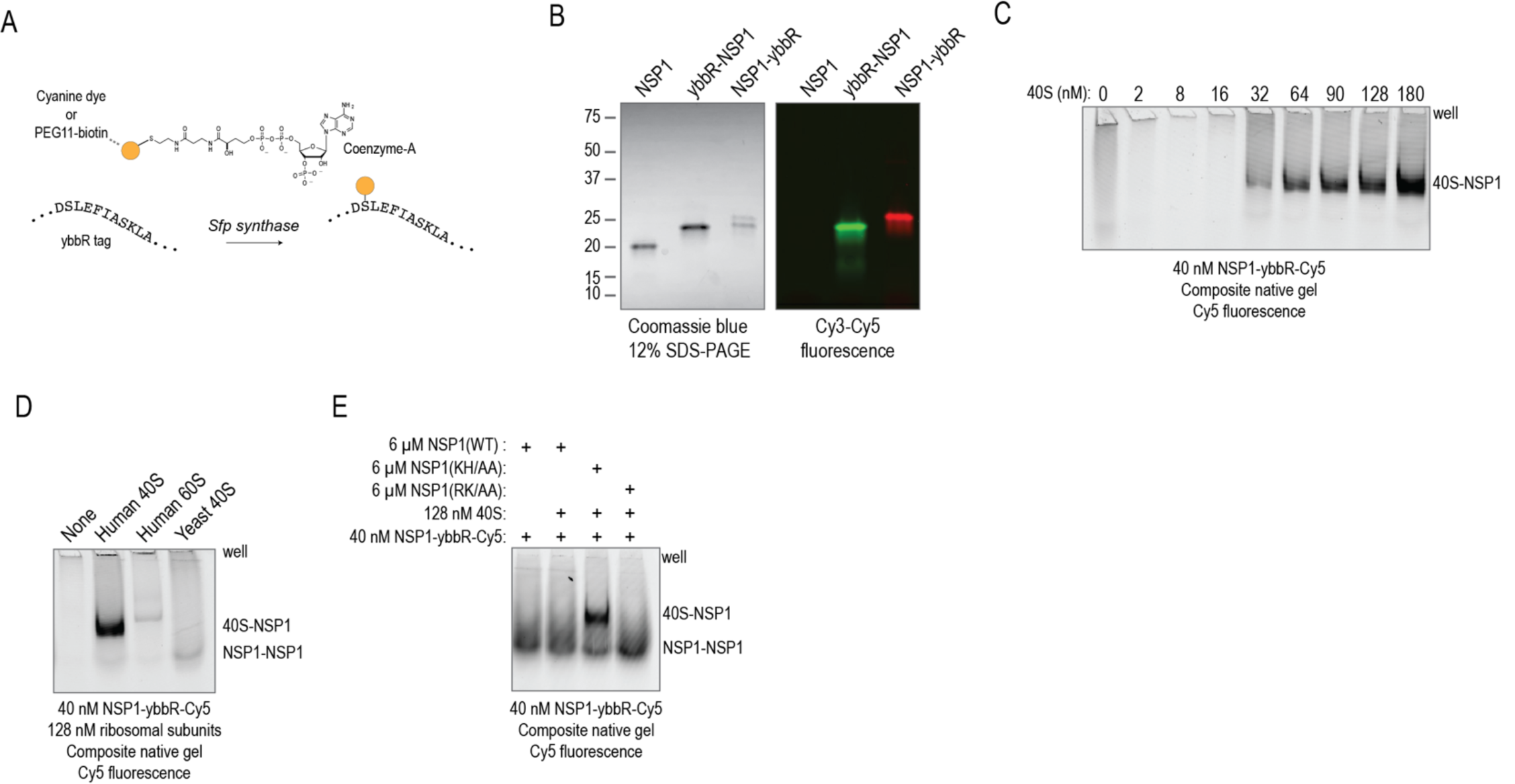
NSP1 specifically bound the human 40S ribosomal subunit. Related to **Fig. 2**. **A)**. Schematic of site-specific, enzymatic attachment of fluorescent cyanine dyes or biotin to the ybbR peptide tag by Sfp synthase. **B)**. Image of purified and fluorescently-labeled NSP1 analyzed via SDS-PAGE. N-terminally tagged ‘ybbR-NSP1’ was labeled with Cy3 fluorescent dye, and C-terminally tagged ‘NSP1-ybbR’ was labeled with Cy5 dye. The image depicts total protein stained with Coomassie blue (left) after scanning the gel for Cy3 and Cy5 fluorescence (right), shown as an overlay of the fluorescent signals. The doublet in the NSP1-ybbR sample likely represents separation of the dye-labeled (upper band) from the non-dye-labeled protein (lower band) in the 12% acrylamide gel. **C)**. Representative fluorescence image of a gel that depicts NSP1-ybbR-Cy5 association with purified and non-labeled human 40S ribosomal subunits at the indicated concentrations, as determined using native gel electrophoresis. NSP1-ybbR-Cy5 was present at 40 nM (by Cy5). Samples were incubated at 37 °C for 10 minutes prior to native gel electrophoresis at 4 °C. *n* = 3. **D)**. Representative fluorescence image of a gel that depicts NSP1-ybbR-Cy5 association with the indicated purified ribosomal subunits, as determined using native gel electrophoresis. 40 nM of NSP1-ybbR-Cy5 was incubated with 128 nM of the indicated subunits at 37 °C for 10 minutes prior to native gel electrophoresis analysis at 4 °C. *n* = 3. **E)**. Representative fluorescence image of a gel that depicts NSP1-ybbR-Cy5 association with purified human 40S ribosomal subunits in the presence of the indicated competitor proteins present at 150-fold molar excess, as determined using native gel electrophoresis at 4 °C. While unlabeled NSP1(WT) and NSP1(RK/AA) eliminated the 40S-NSP1-Cy5 band, substitution of KH to AA in unlabeled NSP1 yielded a strong 40S-NSP1-Cy5 band, similar to experiments in the absence of competitor protein. Thus, NSP1(KH/AA) was unable to associate stably with the human 40S subunit, as expected based on NSP1 from SARS-CoV-1. *n* = 3.

**Supp. Figure 3.**
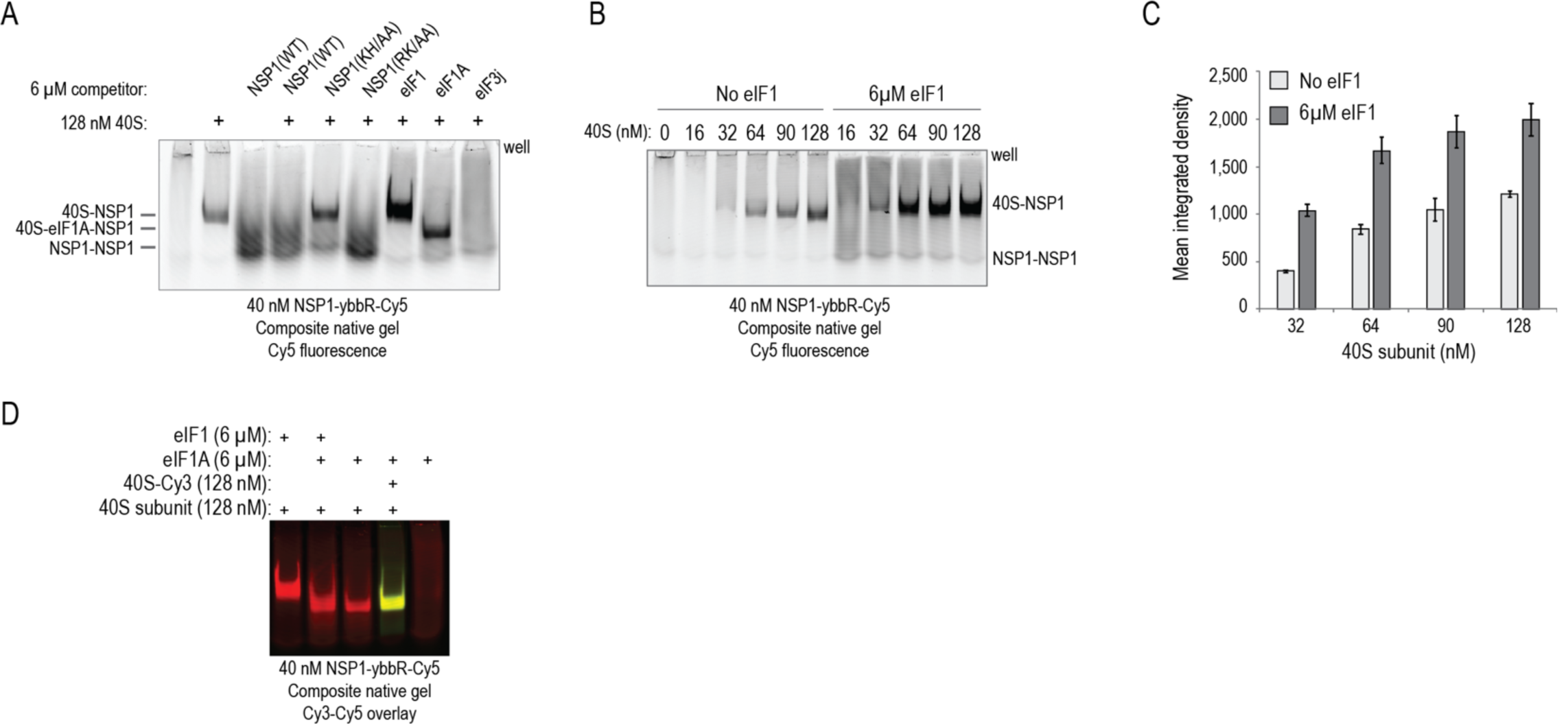
The NSP1–40S subunit interaction was enhanced by eIF1 and blocked by eIF3j. Related to **Fig. 2**. **A)**. Representative fluorescence image of a gel that depicts NSP1-ybbR-Cy5 association with 128 nM purified human 40S ribosomal subunits in the presence of the indicated competitor proteins at 150-fold molar excess, as determined using native gel electrophoresis at 4 °C. All lanes contained 40 nM NSP1-ybbR-Cy5. The identity of the NSP1–eIF1A–40S subunit band was confirmed using fluorescently-labeled 40S ribosomal subunits (panel D). *n* = 3. **B)**. Representative fluorescence image of a gel that depicts eIF1 enhanced NSP1 association with the human 40S ribosomal subunit, as determined using native gel electrophoresis at 4 °C. All lanes contained 40 nM NSP1-ybbR-Cy5. *n* = 3. **C)**. Plot of the mean integrated densities of the indicated bands from the experiment depicted in panel B. At all examined concentrations, the presence of eIF1 increased the intensity of the 40S-NSP1-Cy5 band about 2-fold (mean fold increase = 2 ± 0.4). Error bars represent standard deviation of the mean. See **Supplementary Table 1** for raw data, calculated means, and standard deviations. *n* = 3. **D)**. Representative fluorescence image of a gel that depicts increased migration of the eIF1A–40S subunit complex in composite native gels. All lanes contained 40 nM NSP1-ybbR-Cy5 (red). 40S subunits were labeled with Cy3 (green) via uS19-ybbR (see, Figure 4 and Supp.Fig. 6). The image is an overlay of the Cy3 and Cy5 fluorescence signals. *n* = 3.

**Supp. Figure 4.**
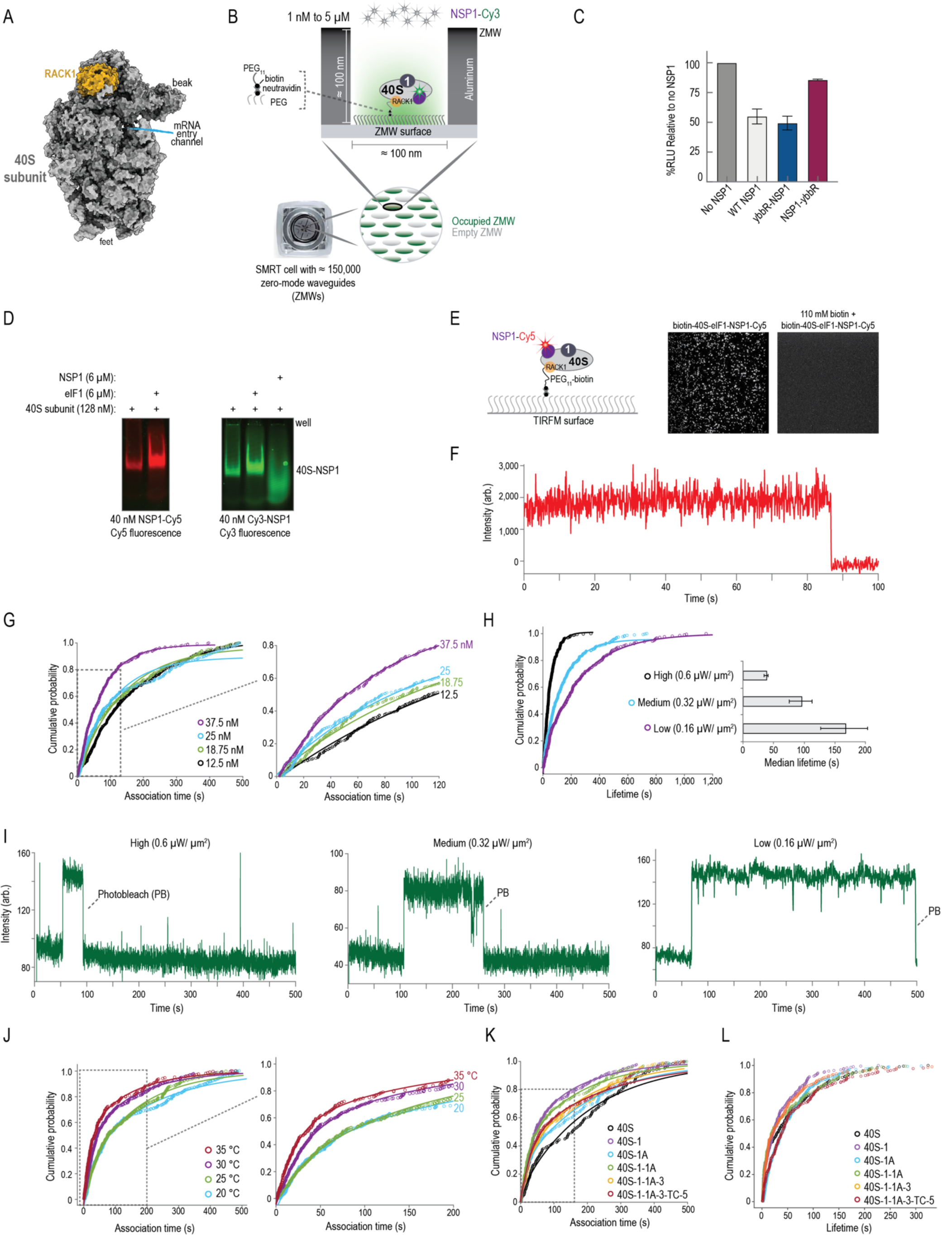
NSP1 bound 40S subunits within pre-initiation complexes. Related to **Fig. 2**. **A)**. Model of the human 40S ribosomal subunit (PDB: 5A2Q). The 40S subunit is depicted in gray and the ribosomal protein RACK1 is in purple. **B)**. Schematic of the zero-mode-waveguide (ZMW) imaging setup. One PacBio SMRT cell contains approximately 150,000 individual ZMWs, which enable direct monitoring of up to four fluorescent signals at concentrations up to 5 µM. The instrument (a PacBio RSII DNA sequencer) enables control of both experimental temperature and the power of the excitation lasers. In this scenario, purified 40S ribosomal subunits that contained RACK1 with a C-terminal ybbR-tag were purified from human cells. In the presence of Sfp synthase, PEG_11_-biotin conjugated to co-enzyme A (co-A) was site-specifically attached to the first serine residue in the 11 amino acid ybbR tag (D**S**LEFIASKLA) on RACK1. Biotinylated 40S subunits (biotin-40S) were incubated with 30-fold molar excess eIF1 at 37 °C for 15 minutes. eIF1–40S subunit complexes were tethered to the neutravidin-coated ZMW surface via the high-affinity biotin–neutravidin interaction. After washing away non-tethered components, data acquisition began at 20 °C and Cy3-NSP1 was added. In this cartoon, ZMWs that contained a Cy3-NSP1 binding event during the experimental time frame are green, and ZMWs that lacked a binding event are gray. **C)**. Translation inhibition activity of the labeled proteins. Plot of mean nLuc signal from IVT reactions programed with GAPDH reporter mRNA in the absence of NSP1 or the presence of 0.4 μM of WT, N-terminally (ybbR-NSP1), or C-terminally tagged (NSP1-ybbR) NSP1. Error bars represent SEM. *n = 3*. **D)**. Representative fluorescence image of a gel that compares N-terminally (ybbR-NSP1) and C-terminally tagged (NSP1-ybbR) NSP1 association with purified human 40S ribosomal subunits. The Cy5 (left) and Cy3 (right) images were run in parallel on the same gel, but they were separated here for clarity. Wild-type, unlabeled NSP1 or eIF1 was added at 6 µM. 40 nM of labeled NSP1 (either Cy5-labeled or Cy3-labeled) was present. *n* = 3. **E)**. Schematic of total internal reflection fluorescence microscopy (TIRFM) experiments performed at equilibrium. Biotinylated 40S (biotin-40S) subunits were incubated with 12-fold molar excess of NSP1-Cy5 and 150-fold excess eIF1 at 37 °C for 15 minutes. Pre-formed complexes were incubated on a TIRFM surface coated with neutravidin. After washing away unbound complexes and components, we imaged tethered ribosomal complexes at room temperature in the presence of 1 µM eIF1 and 4 nM of NSP1-Cy5. We observed a few hundred NSP1-Cy5 molecules tethered to the imaging surface in all view fields examined (representative view field shown, *n* = 18). Consistent with specific tethering, NSP1-Cy5 signal was eliminated by pre-saturation of the imaging surface with 110 mM biotin. **F)**. Example single-molecule fluorescence trace from the left view field of the equilibrium TRIFM experiment outlined in panel E (no biotin pre-treatment). The trace begins with fluorescence signal from NSP1-Cy5 (red) bound to a tethered eIF1–40S subunit complex. **G)**. Plot of the cumulative probability of Cy3-NSP1 association times at the indicated concentrations with tethered eIF1–40S subunit complexes at 20 °C. The left plot depicts all association times, and the right plot highlights the subset within the dashed box of the left plot. Lines represent fits to single-exponential functions. See **Supplementary Table 2** for samples sizes and the parameters for relevant fits. **H)**. Plot of the cumulative probability of (left) and median (right) Cy3-NSP1 lifetimes on eIF1–40S subunit complexes at 20 °C and the indicated excitation laser powers. Cy3-NSP1 was added at 25 nM. The wavelength of the excitation laser was 532 nm, and it was used at high (0.6 µW/ µm^2^), medium (0.32 µW/ µm^2^), and low (0.16 µW/ µm^2^) power to determine whether the lifetime of the NSP1 fluorescence signal was limited by the photostability of the Cy3 dye. Lines represent (left plot) fits to exponential functions. Error bars (right plot) represent 95% C.I. of the median values. See panel I for example traces. See **Supplementary Table 2** for samples sizes and the parameters for relevant fits. **I)**. Example single-molecule fluorescence trace that depicts association of Cy3-NSP1 (green) with a tethered eIF1–40S subunit complex acquired at different laser powers. The wavelength of the excitation laser was 532 nm, and it was used at high (0.6 µW/ µm^2^), medium (0.32 µW/ µm^2^), and low (0.16 µW/ µm^2^) power to determine whether the lifetime of the Cy3-NSP1 fluorescence signal was limited by the photostability of the Cy3 dye. During imaging, eIF1 was present at 1 µM. Loss of fluorescence due to likely photobleaching of the Cy3 dye is indicated on each trace. **J)**. Plot of the cumulative probability of Cy3-NSP1 association times with tethered eIF1–40S subunit complexes at the indicated temperatures. In all experiments, Cy3-NSP1 was present at 25 nM. The left plot depicts all association times, and the right plot highlights the subset within the dashed box of the left plot. Lines represent fits to double-exponential functions. See **Supplementary Table 2** for samples sizes and the parameters for relevant fits. **K)**. Plot of the cumulative probability of Cy3-NSP1 association times with the indicated ribosomal pre-initiation complexes. In all experiments, Cy3-NSP1 was present at 25 nM and the temperature was 30 °C. The dashed box represents the region of the plot displayed in **Fig. 2F**. Lines represent fits to double-exponential functions. See **Supplementary Table 2** for samples sizes and the parameters for relevant fits. **L)**. Plot of the cumulative probability of Cy3-NSP1 lifetimes on the indicated 40S pre-initiation complexes. In all experiments, Cy3-NSP1 was present at 25 nM and the temperature was 30 °C. See **Supplementary Table 2** for samples sizes and medians with 95% C.I..

**Supp. Figure 5.**
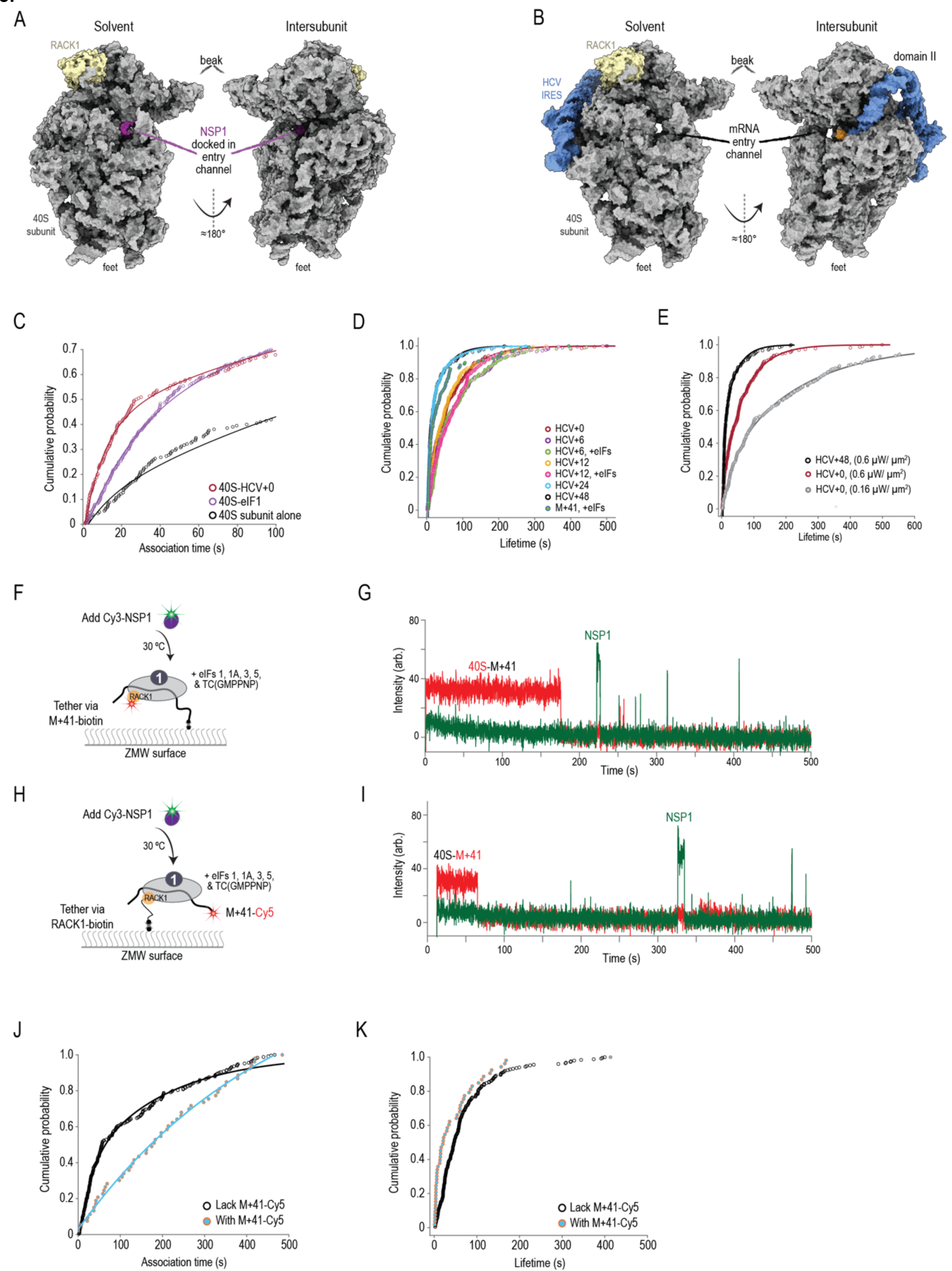
mRNA within the mRNA entry channel of the 40S subunit inhibited NSP1 association. Related to **Fig. 3**. **A)**. Model of NSP1 bound to the mRNA entry channel of the human 40S subunit (PDB:6ZLW). The 40S subunit is depicted in gray, RACK1 in light yellow, and NSP1 in purple. **B)**. Model of the human 40S subunit bound by the HCV IRES (PDB: 5A2Q). The 40S subunit is depicted in gray, RACK1 in light yellow, and the HCV IRES in blue. The model of the IRES ends at the start codon (AUG, highlighted in orange), leaving the mRNA entry channel of the 40S subunit empty. Domain II of the IRES holds the head of the 40S subunit in the open conformation. **C)**. Plot of the cumulative probability of Cy3-NSP1 association times with 40S subunit complexes. In all experiments, Cy3-NSP1 was present at 25 nM and the temperature was 30 °C. The data for the 40S subunit alone (40S) and the eIF1–40S subunit complex (40S– eIF1) were replotted here from **Fig. 2F** and **Supp.Fig. 4K** to provide easy comparison to the data for NSP1 association with the 40S– HCV+0 complex. Lines represent fits to single-or double-exponential functions. See **Supplementary Table 3** for samples sizes and the parameters for relevant fits. **D)**. Cumulative probability plot of observed lifetimes of 25 nM Cy3-NSP1 with the indicated 40S–mRNA complexes at 30 °C. ‘+eIFs’ indicates eIFs 1, 1A, 3, 5, and TC(GMPPNP) were included at all stages of the experiment. Lines represent fits to exponential functions. See **Supplementary Table 3** for samples sizes, medians with 95% C.I., and parameters for fits. **E)**. Cumulative probability plot of observed lifetimes of 25 nM Cy3-NSP1 with the indicated 40S–mRNA complexes at 30 °C. The power of the excitation laser (532 nm) is indicated (high power = 0.6 µW/µm^2^, low power = 0.16 µW/µm^2^). HCV+48 and HCV+0 data at high laser power were replotted here from panel D for presentation purposes. Lines represent fits to double-exponential function. See **Supplementary Table 3** for samples sizes, medians with 95% C.I., and parameters for fits. **F)**. Schematic of the single-molecule fluorescence experimental setup with biotinylated M+41 model mRNA. 40S ribosomal subunits were labeled with Cy5 dye via RACK1-ybbR. M+41 RNA was labeled on the 3’-terminus with biotin. Pre-formed M+41–40S-Cy5 complexes were tethered to the ZMW imaging surface in the presence of eIF1, eIF1A, eIF3, eIF5, and TC-GMPPNP. At the start of data acquisition, Cy3-NSP1 was added at 25 nM at 30 °C. **G)**. Example single-molecule fluorescence trace that depicts a tethered M+41–40S-Cy5 complex and subsequent association of Cy3-NSP1. The 40S subunit and ybbR-NSP1 were labeled with Cy5 (red) and Cy3 (green) dyes, respectively. Raw fluorescence intensities were corrected in this image to set baseline intensities to zero for presentation. The association time (Δt) was defined as time elapsed from the addition of Cy3-NSP1 until the burst of Cy3 fluorescence (green), which signified NSP1 association. The lifetime was defined as the duration of the Cy3 fluorescence signal. **H)**. Schematic of the single-molecule fluorescence experimental setup with fluorescently-labeled M+41 model mRNA. 40S ribosomal subunits were labeled with biotin via RACK1-ybbR. M+41 RNA was labeled on the 3’-terminus with Cy5. Pre-formed Cy5-M+41–40S subunit complexes were tethered to the ZMW imaging surface in the presence of eIF1, eIF1A, eIF3, eIF5, and TC-GMPPNP. At the start of data acquisition, Cy3-NSP1 was added at 25 nM at 30 °C. **I)**. Example single-molecule fluorescence trace that depicts a tethered Cy5-M+41–40S subunit complex and subsequent association of Cy3-NSP1. The M+41 RNA and ybbR-NSP1 were labeled with Cy5 (red) and Cy3 (green) dyes, respectively. Raw fluorescence intensities were corrected in this image to set baseline intensities to zero for presentation. The association time (Δt) was defined as time elapsed from the addition of Cy3-NSP1 until the burst of Cy3 fluorescence (green), which signified NSP1 association. The lifetime was defined as the duration of the Cy3 fluorescence signal. **J, K)**. Cumulative probability plots of the observed association times (panel J) and lifetimes (panel K) of Cy3-NSP1 with the indicated 40S–mRNA complexes. Biotinylated 40S subunits (as in **Fig. 2** & **Supp.Fig. 4**) were directly tethered to the surface in the presence of the model mRNA (M+41) labeled on the 3’-terminus with Cy5 and eIFs 1,1A, 5, and TC-GMPPNP. Both sets of molecules plotted were here were obtained from the same experiment. The ‘Lack M+41’ data represent ZMWs that had at least one Cy3-NSP1 binding event but lacked Cy5-M+41 signal. The ‘With M+41’ data represent ZMWs that began with Cy5 signal (Cy5-M+41) and contained at least one NSP1 binding event (Cy3) longer than ≈ 5s. Cy3-NSP1 was present at 25 nM and the temperature was 30 °C. See **Supplementary Table 3** for samples sizes, parameters for relevant fits, and medians with 95% C.I..

**Supp. Figure 6.**
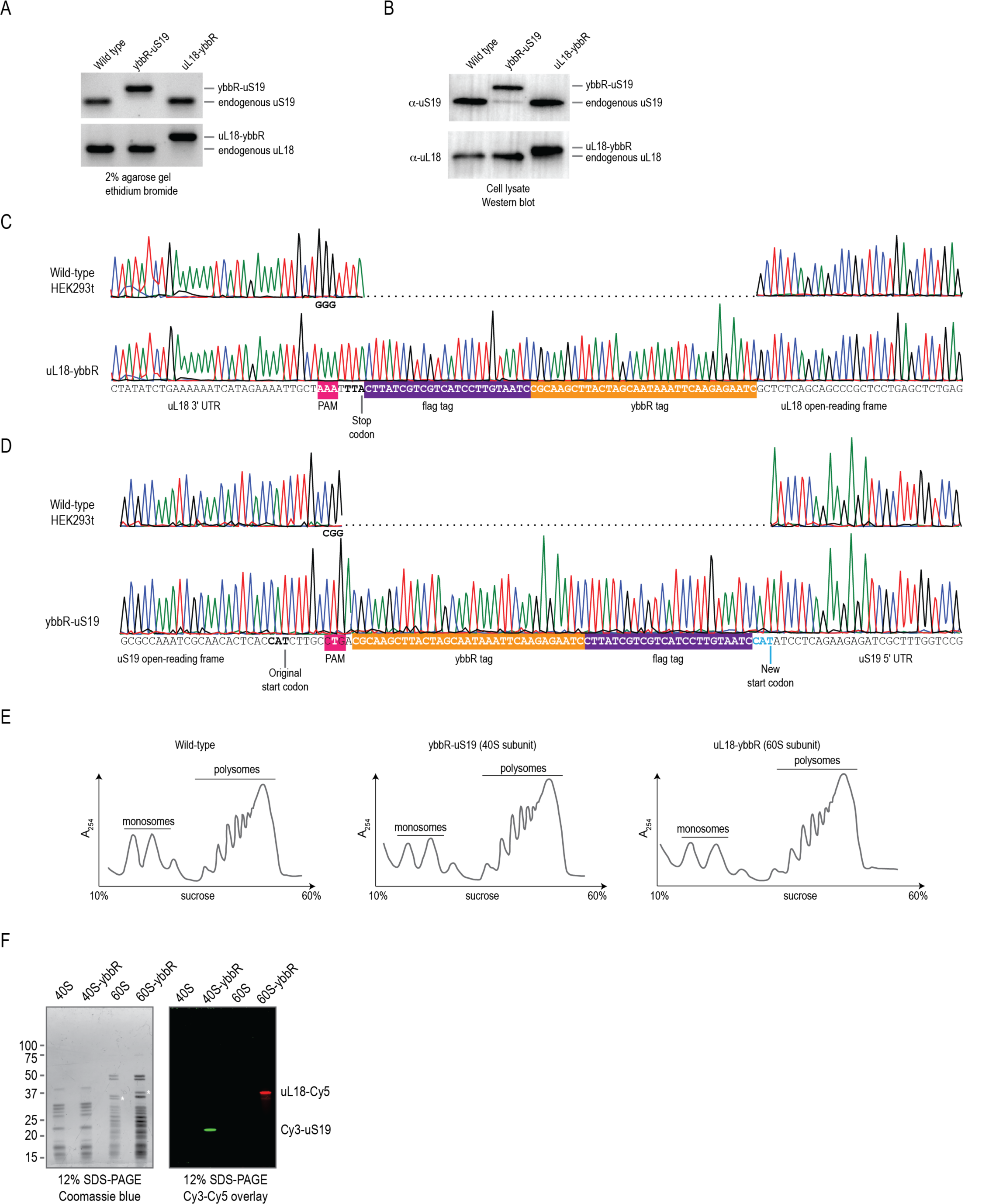
Generation of an inter-subunit FRET signal for the human ribosome. Related to **Fig. 4**. **A)**. Image of a 2% agarose gel stained with ethidium bromide. The analyzed PCR products were generated with primers that flanked the desired insertion site for the ybbR-flag tag, with at least one of them lying outside the repair template. **B)**. Image of Western blots using the indicated antibodies on whole cell extract from the indicated cell lines. Together with the PCR data from panel B, these data indicated that all endogenous copies of uS19 or uL18 were edited to incorporate ybbR-flag tags. The faint band that may represent endogenous uS19 in the ybbR-uS19 sample of the Western blot could result from translation initiation at the original start codon, rather than the new, inserted start codon. **C, D)**. Traces from Sanger sequencing of PCR products generated as in panel B for the indicated cell lines and genomic loci. Important regions within the sequenced DNA segments are annotated. For both ybbR-uS19 and uL18-ybbR cell lines, the ybbR-flag tag was inserted as designed and the used PAM sites were disrupted, as encoded in the repair templates. **E)**. Representative traces from polysome profiling assays on the indicated cell lines. The positions of monosomes (individual ribosomal subunits and single 80S ribosomes) and polysomes (multiple 80S ribosomes) are indicated. Similar levels of polysomes were detected in wild-type, ybbR-uS19, and uL18-ybbR HEK293t cell lines. Thus, ribosomes that contain the ybbR-flag tag on uS19 or uL18 were functional. *n = 2*. **F)**. Gel that depicts purified and fluorescently-labeled ribosomal subunits. Ribosomal subunits were purified from either the HEK293T cell line that expressed ybbR-uS19 (40S-ybbR & 60S samples) or uL18-ybbR (40S & 60S-ybbR samples). Following ybbR-labeling reactions with either Cy3 (green) or Cy5 (red) dyes, ribosomal subunits were analyzed by SDS-PAGE on 12% acrylamide gels. Prior to staining total protein with Coomassie blue (left image), gels were scanned for both Cy3 and Cy5 fluorescence (right image), which is presented as an overlay of both fluorescent signals.

**Supp. Figure 7.**
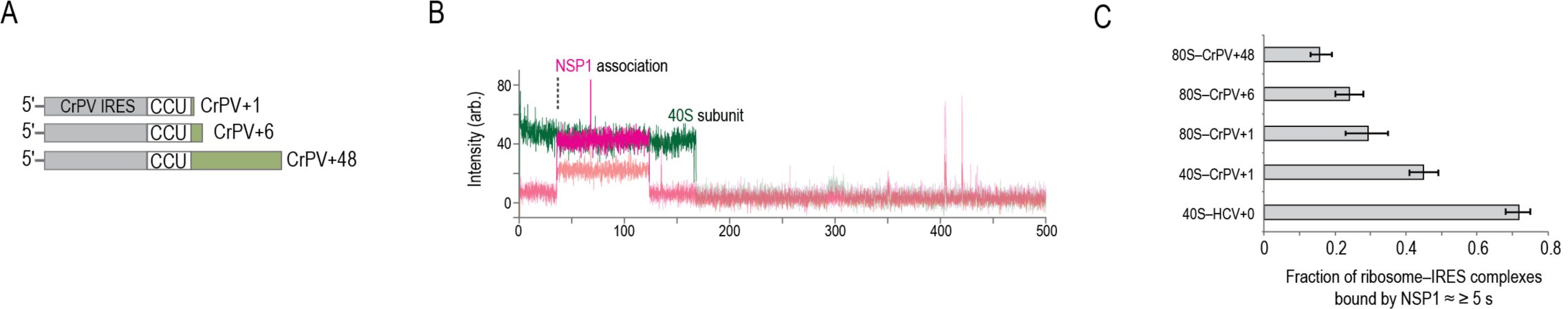
NSP1 inefficiently bound to 80S–CrPV IRES complexes. Related to **Fig. 4**. **A)**. Schematic of CrPV IRES RNAs used in the assays. The RNAs contained 1, 6, or 48 nucleotides downstream of the CCU codon in the CrPV IRES, which is present in the ribosomal A site. All RNAs were biotinylated on the 5’-terminus. **B)**. Example single-molecule fluorescence trace that depicts addition of Cy5.5-NSP1 to tethered 40S–CrPV+1 complexes. The 40S subunit was labeled with Cy3 (green) and NSP1 with Cy5.5 (magenta). Due to bleed through across the three fluorescent channels, the Cy3, Cy5, and Cy5.5 signals were made transparent before and after relevant events for presentation here. The association time (Δt) was defined as the time elapsed from the addition of Cy5.5-NSP1 until the burst of Cy5.5 fluorescence (magenta), which signified NSP1 association. The lifetime was defined as the duration of the Cy5.5 fluorescence signal. **C)**. Plot of the fraction of the indicated ribosome–IRES complexes bound at least once by NSP1 for ≥ ≈ 5s. Error bars represent 95% C.I. yielded from bootstrapping analyses.

**Supp. Figure 8.**
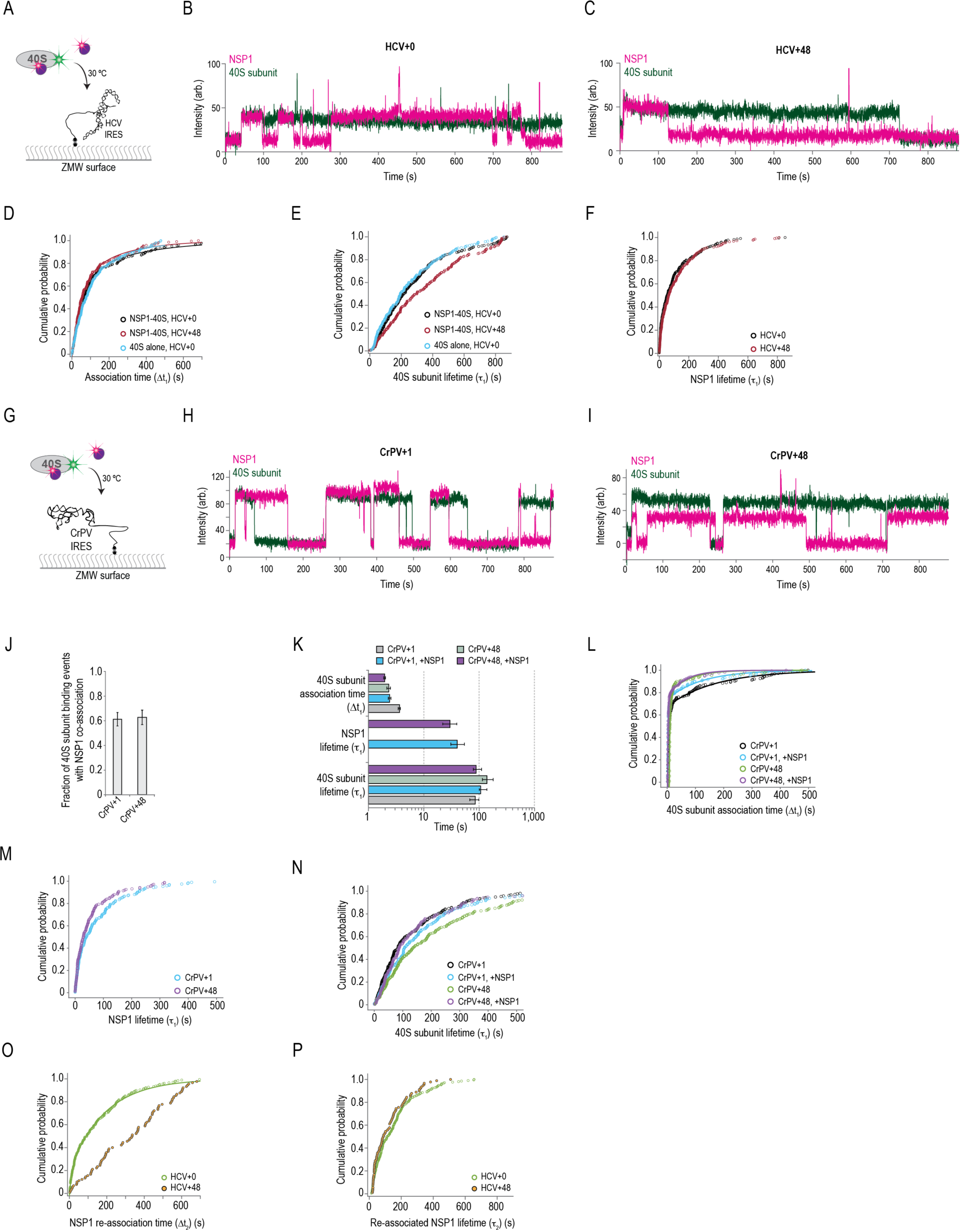
NSP1 remained bound to 40S subunits upon association with model mRNAs. Related to **Fig. 5**. **A)**. Schematic of the single-molecule fluorescence experimental setup with the HCV IRES. 40S ribosomal subunits were labeled with Cy3 dye via uS19-ybbR. When indicated, NSP1–40S subunit complexes were pre-formed by incubating 2-fold molar excess Cy5.5-NSP1 with 40S-Cy3 subunits at 37 °C for 15 minutes. After tethering of HCV+0 or HCV+48 model RNAs in ZMWs and start of data acquisition, a final concentration of 15 nM (by 40S subunits) 40S-Cy3 subunits or NSP1–40S subunit complexes were added at 30 °C. **B**,**C)**. Example single-molecule fluorescence traces that depict association of NSP1–40S subunit complexes with tethered HCV+0 (panel B) or HCV+48 (panel C) IRES molecules. The 40S subunit and NSP1 were labeled with Cy3 (green) and Cy5.5 (magenta) dyes, respectively. Raw fluorescence intensities were corrected in this image to set baseline intensities to zero for presentation. These traces depict the full duration of data acquisition. In both, a single NSP1–40S subunit complex associates with the HCV IRES. Following initial loss of NSP1 signal, NSP1 readily re-associated with the same 40S–HCV+0 complex but not the 40S–HCV+48 complex. **D)**. Cumulative probability plot of observed association times of either 40S subunits alone or NSP1–40S subunit complexes with the indicated model RNAs at 30 °C. Lines represent fits to double-exponential functions. See **Supplementary Table 5** for samples sizes and the parameters for relevant fits. **E**,**F)**. Cumulative probability plots of 40S subunit (panel E) and NSP1 (panel F) lifetimes on the indicated model RNAs at 30 °C. See **Supplementary Table 5** for samples sizes and medians with 95% C.I.. **G)**. Schematic of the single-molecule fluorescence experimental setup with the CrPV IRES. 40S ribosomal subunits were labeled with Cy3 dye via uS19-ybbR. When indicated, NSP1–40S subunit complexes were pre-formed by incubating 2-fold molar excess Cy5.5-NSP1 with 40S-Cy3 subunits at 37 °C for 15 minutes. After tethering of CrPV+1 or CrPV+48 model RNAs in ZMWs and start of data acquisition, a final concentration of 15 nM (by 40S subunits) 40S-Cy3 subunits or NSP1–40S subunit complexes were added at 30 °C. **H**,**I)**. Example single-molecule fluorescence traces that depict association of NSP1–40S subunit complexes with a tethered CrPV+1 (panel H) or CrPV+48 (panel I) IRES molecules. The 40S subunit and NSP1 were labeled with Cy3 (green) and Cy5.5 (magenta) dyes, respectively. Raw fluorescence intensities were corrected in this image to set baseline intensities to zero for presentation. These traces depict the full duration of data acquisition. The 40S–CrPV IRES complex was more transient than that of the 40S–HCV IRES complex. Thus, we often observed multiple NSP1–40S subunit complex association events within a single ZMW, particularly for CrPV+1, which prevented analysis of NSP1 re-association with these model RNAs. **J)**. Plot of the fraction of 40S subunit binding events with NSP1 co-association on the indicated model RNAs at 30 °C. Error bars represent the 99% C.I.. **K)**. Plot of association times and median lifetimes from the indicated experiments using CrPV+1 or CrPV+48 model RNAs at 30 °C. ‘+NSP1’ indicates experiments where pre-formed NSP1–40S subunit complexes were added. The association time (40S Δt_1_) was defined as the reciprocal of the fast association rate derived from fits to double-exponential functions, with error bars indicating the 95% C.I. of the rate. Error bars for median lifetimes (40S *τ*1, NSP1 *τ*_1_) represent 95% C.I.. Due to the relative instability of these IRES-40S subunit complexes and the rapid re-association of 40S subunits, NSP1 re-association with single IRES–40S subunit complexes was not examined. See **Supplementary Table 5** for samples sizes and the parameters for relevant fits. **L-N)**. Cumulative probability plots of the indicated association times (panel L) or lifetimes (panels M, N) in experiments with CrPV+1 or CrPV+48 model RNAs at 30 °C. ‘+NSP1’ indicates experiments where pre-formed NSP1–40S subunit complexes were added. Lines represent fits to double-exponential functions. See **Supplementary Table 5** for samples sizes, parameters for relevant fits, and medians with 95% C.I.. **O**,**P)**. Cumulative probability plots of observed NSP1 re-association times (panel O) or lifetimes (panel P) with the indicated IRES–40S subunit complexes at 30 °C. As outlined in **Fig. 4A**, we focused on ZMWs where a single 40S subunit associated within the first 200 s (≈ 75% of all events, see panel D). We then quantified the time elapsed from the loss of the first Cy5.5 signal to the next burst of Cy5.5 fluorescence at least ≈ 20 s in length (≈ 70% of initial NSP1 binding events, see panel F), which was defined as the NSP1 re-association time (NSP1 Δt_2_). The duration of this second Cy5.5 event was defined as the re-associated NSP1 lifetime (NSP1 *τ*_2_). Lines represent fits to double-exponential functions. See **Supplementary Table 5** for samples sizes, parameters for relevant fits, and medians with 95% C.I..

